# Ras-mediated homeostatic control of front-back signaling dictates cell polarity

**DOI:** 10.1101/2023.08.30.555648

**Authors:** Yiyan Lin, Dhiman Sankar Pal, Parijat Banerjee, Tatsat Banerjee, Guanghui Qin, Yu Deng, Jane Borleis, Pablo A. Iglesias, Peter N. Devreotes

**Affiliations:** Department of Cell Biology and Center for Cell Dynamics, School of Medicine, Johns Hopkins University, Baltimore, MD, USA; Department of Biological Chemistry, School of Medicine, Johns Hopkins University, Baltimore, MD, USA; Department of Physics & Astronomy, Johns Hopkins University, Baltimore, MD, USA; Department of Chemical and Biomolecular Engineering, Whiting School of Engineering, Johns Hopkins University, Baltimore, MD, USA; Department of Computer Science, Whiting School of Engineering, Johns Hopkins University, Baltimore, MD, USA; Department of Electrical and Computer Engineering, Whiting School of Engineering, Johns Hopkins University, Baltimore, MD, USA

**Keywords:** cancer metastasis, biochemical excitability, actin cytoskeleton, optogenetics, signaling

## Abstract

Studies in the model systems, *Dictyostelium* amoebae and HL-60 neutrophils, have shown that local Ras activity directly regulates cell motility or polarity. Localized Ras activation on the membrane is spatiotemporally regulated by its activators, RasGEFs, and inhibitors, RasGAPs, which might be expected to create a stable ‘front’ and ‘back’, respectively, in migrating cells. Focusing on C2GAPB in amoebae and RASAL3 in neutrophils, we investigated how Ras activity along the cortex controls polarity. Since existing gene knockout and overexpression studies can be circumvented, we chose optogenetic approaches to assess the immediate, local effects of these Ras regulators on the cell cortex. In both cellular systems, optically targeting the respective RasGAPs to the cell front extinguished existing protrusions and changed the direction of migration, as might be expected. However, when the expression of C2GAPB was induced globally, amoebae polarized within hours. Furthermore, within minutes of globally recruiting either C2GAPB in amoebae or RASAL3 in neutrophils, each cell type polarized and moved more rapidly. Targeting the RasGAPs to the cell backs exaggerated these effects on migration and polarity. Overall, in both cell types, RasGAP-mediated polarization was brought about by increased actomyosin contractility at the back and sustained, localized F-actin polymerization at the front. These experimental results were accurately captured by computational simulations in which Ras levels control front and back feedback loops. The discovery that context-dependent Ras activity on the cell cortex has counterintuitive, unanticipated effects on cell polarity can have important implications for future drug-design strategies targeting oncogenic Ras.

## Introduction

Ras GTPases play a vital role in transmitting signals within cells, influencing growth and survival, and activating mutations in Ras genes are found in ∼30% of all cancers^1–3^. As a result, targeting mutant Ras has become a major focus in cancer drug development. While extensive research has been dedicated to understanding the role of Ras mutations in promoting cancer growth, the impact of these mutations on cell migration and metastasis has received relatively less attention. Nevertheless, studies conducted in the model systems, *Dictyostelium* amoebae and human neutrophils, suggest that local Ras activity plays a direct and immediate role in cell polarity and motility^4–9^.

Biochemical and genetic investigations have revealed that Ras is activated both by stimulation of G-protein coupled receptors and spontaneously at protrusions^6,10–13^. Ras function is spatiotemporally controlled by its activators, RasGEFs, and inhibitors, RasGAPs, which might be expected to create ’front’ and ’back’, respectively, during cell migration^11,14–19^. However, Ras activity must be carefully balanced, as constitutively active Ras expression can lead to hyper-activation of the PI3K/TORC2-PKB pathways, resulting in significant cell spreading and defective migration^5,20,21^. Additionally, knockout/knockdown studies of RasGAPs, such as NF1, C2GAP1, C2GAP2, and CAPRI have demonstrated their significant impact on Ras and protrusive activities^8,11,17,18,22,23^. For example, knockout of NF1 in *Dictyostelium* causes increased macropinocytosis, and moreover, human genetic deletion of NF1 results in severe neurofibromatosis type 1^24–26^.

Understanding the physiological relevance of manipulating Ras activity is of paramount importance, but knockout studies of Ras proteins have proven relatively ineffective, possibly due to redundancy. For example, *Dictyostelium* cells lacking Ras isoforms continue to grow and directed migration remains little affected^27–29^. Furthermore, while new treatments with small molecule inhibitors targeting constitutively active KRasG12C show promise, they also pose significant challenges^30–32^. A potentially powerful alternative route of inhibiting Ras activity would be to activate the RasGAP proteins. Indeed, a recent study showed that locally activating a RasGAP can inhibit Ras and halt random and directed migration in immune cells^4^.

We designed a series of studies to test the capacity of RasGAPs to manipulate Ras activity. Two different RasGAP proteins, C2GAPB and RASAL3, were able to inhibit Ras activity in *Dictyostelium* and differentiated HL-60 neutrophils, respectively, but surprisingly, under certain conditions, these cells become highly polarized and migrate rapidly. Our evidence suggests that the polarization was due to an increase in suppression of Ras activity leading to increased contraction at the back. Taken together with previous observations of hyper-activated Ras activity, our study shows that there is not a direct linear relationship between Ras activation and migration, and an optimal level of activated Ras leads to polarity and migration. Our study has important implications for interventions, such as small molecule inhibitors, designed to simply reduce Ras activity as a reduction could cause unanticipated effects such as exaggerated metastasis.

## Results

### RasGAPs inhibit Ras and PI3K activities to shut off cellular protrusions

To investigate the impact of Ras activity on cell migration and polarity, we examined the effects of RasGAP, C2GAPB, in modulating Ras activity and downstream PI3K activity. Since we encountered challenges in expressing RasGAP proteins using standard expression methods, we developed a doxycycline-inducible system to improve C2GAPB expression. This system not only facilitated efficient expression of RasGAP proteins but also allowed for convenient comparison of results in the absence or presence of RasGAP expression. We first compared Ras or PI3K activities by expressing fluorescence-labeled biosensors, RBD or PHcrac, respectively, with or without C2GAPB expression. Confocal imaging of the midline of cells revealed broad patches of RBD and PIP3 at the protrusions in the absence of C2GAPB. However, upon overnight induction of C2GAPB expression, these patches reduced in size (Figure 1A-C). Previous studies have shown that the Ras/PIP3 patches underlie protrusions which mediate cell movement and macropinocytosis^11,33,34^. Uptake measurements showed that macropinocytosis was significantly reduced upon C2GAPB expression (Figure S1A and B).

**Figure 1.**
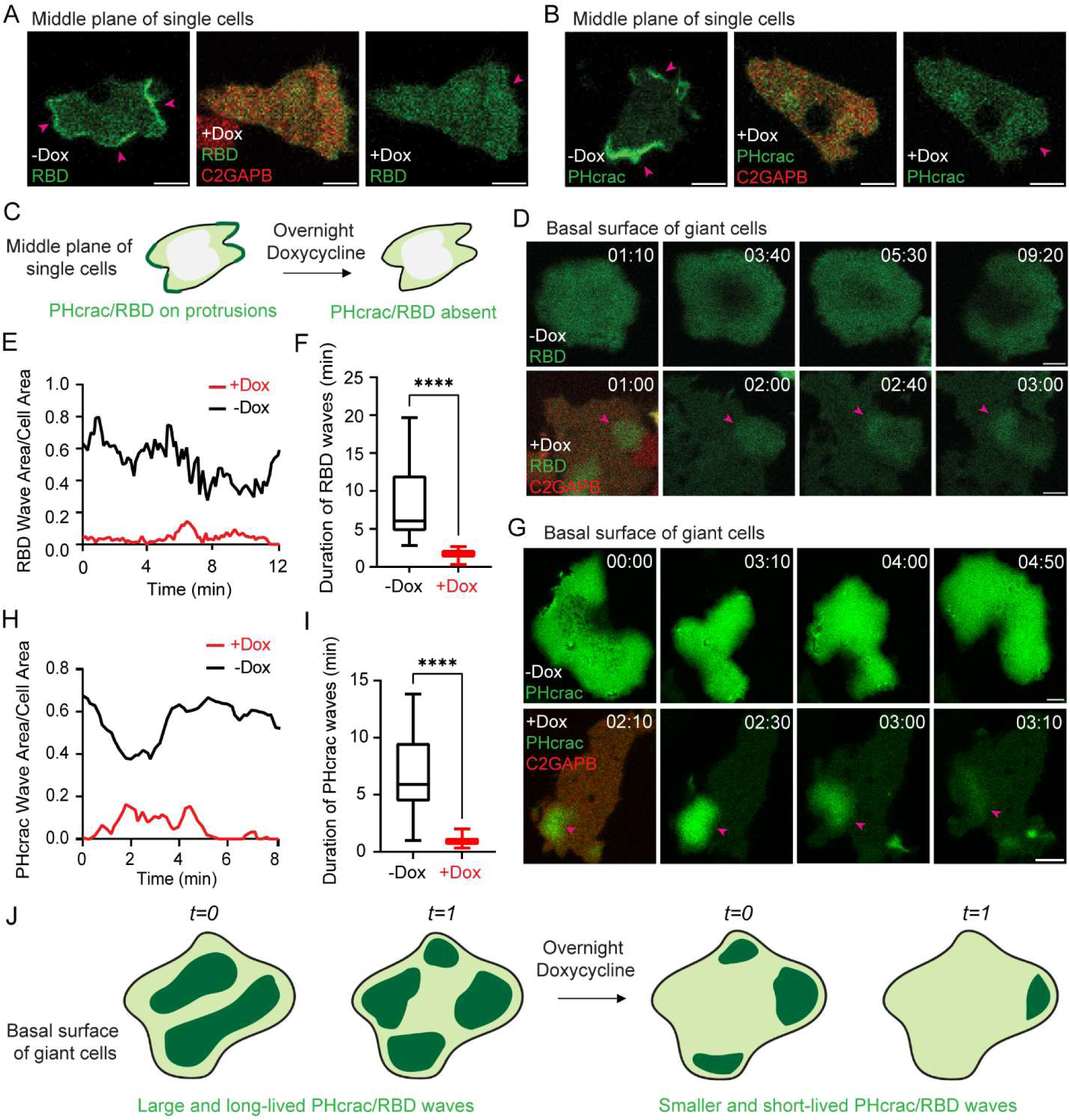
C2GAPB inhibits Ras and PI3K activities in single and electrofused *Dictyostelium*. Confocal images of vegetative *Dictyostelium* single cells expressing **(A)** GFP-RBD (biosensor for activated Ras; green) or **(B)** PHcrac-YFP (biosensor for PIP3; green) before and after doxycycline-induced mRFPmars-C2GAPB (red) expression. Pink arrows highlight fronts of these cells. Scale bars represent 5 µm. **(C)** Cartoon summarizes our observations in (A) and (B) that both Ras activation and PIP3 level on the cell membrane significantly reduce with C2GAPB expression. Time-lapse confocal images of vegetative *Dictyostelium* electrofused or ‘giant’ cells expressing **(D)** GFP-RBD (green) or **(G)** PHcrac-YFP (green) before and after doxycycline-induced mRFPmars-C2GAPB (red) expression. Pink arrows point at reduced RBD or PHcrac waves in presence of C2GAPB. Time in min:sec format. Scale bars represent 5 µm. RBD or PHcrac wave area **(E or H)** and duration **(F or I)** before (‘-Dox’; black) and after (‘+Dox’; red) doxycycline-induced mRFPmars-C2GAPB expression. **(J)** Cartoon depicting reduction in size and duration of RBD and PHcrac waves on the basal surface of giant *Dictyostelium* cells, after overnight doxycycline induction of C2GAPB.

Activities of Ras and PI3K have been extensively documented to form broad, propagating waves on the basal surface of electrofused, giant *Dictyostelium* cells^12,13,35–40^. Thus, we further investigated the effects of C2GAPB on Ras/PIP3 waves and found that both RBD and PHcrac waves became smaller and more transient upon expressing C2GAPB (Figure 1D and G; Videos S1-4). Quantitative analysis revealed that in the absence of C2GAPB expression, the area covered by RBD or PHcrac waves at steady-state varied between 35% and 80% of the total cell area, but with induction of C2GAPB, the wave area decreased to less than 20% (Figure 1E and H). The mean duration of a wave also decreased from 5 minutes to less than a minute (Figure 1F and I). Therefore, inducible C2GAPB expression led to a significant reduction of RBD and PHcrac patches and waves (Figure 1C and J). Conversely, in C2GAPB null cells, wide, propagating RBD waves were observed, validating that C2GAPB expression suppresses Ras activity (Figure S1C)^11^.

To further investigate the functional implications of C2GAPB on cellular protrusions and cytoskeleton remodeling, we utilized optogenetics to transiently recruit C2GAPB to the plasma membrane. Based on our observations that activated Ras and PIP3 levels decreased with C2GAPB expression, we hypothesized that the recruitment of C2GAPB could inhibit cellular protrusions. In our system, upon 488 nm laser stimulation, mRFPmars-SspB fused C2GAPB is recruited to membrane anchor N150-ilID (Figure 2A). When C2GAPB was recruited to the cellular protrusions at the leading edge (indicated with white arrows), mature protrusions quickly vanished causing the cell to contract, while new protrusions emerged at the former back end of the cell (shown with pink arrows). Consequently, the cell moved in the opposite direction (Figure 2A and C). Angular histogram analyses revealed that the probability of nascent protrusion formation was highest at ∼120-150 degrees away from the C2GAPB recruitment area (Figure 2D). In contrast, although there was a slight effect of light, recruitment of the empty control vector did not block the production of new protrusions, and the cell continued to move close to its original direction (Figure 2E). Angular histogram showed that the highest probability of new protrusion formation occurred at ∼0-40 degrees from the C2GAPB recruitment area (Figure 2F).

**Figure 2.**
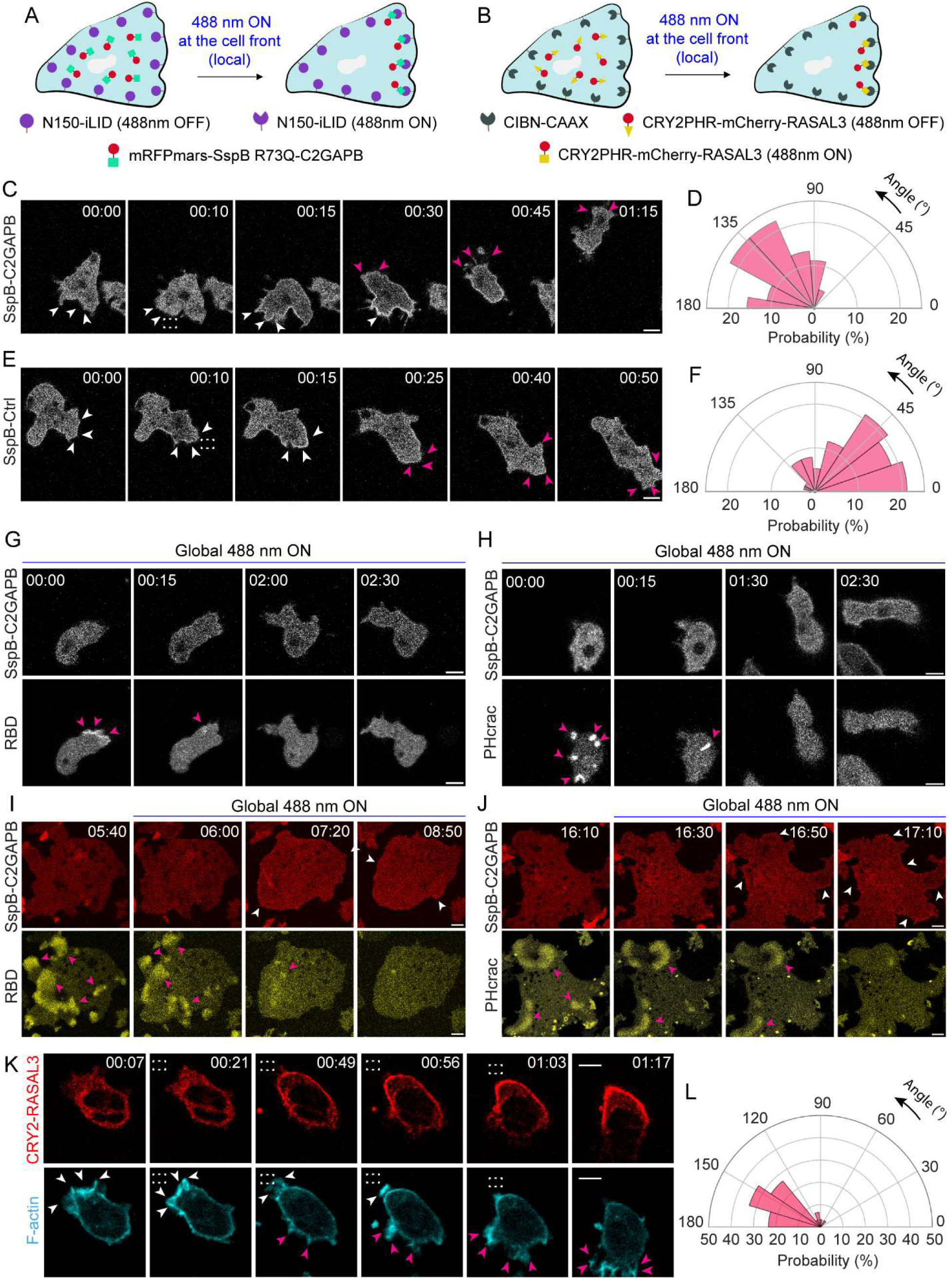
Recruitment of C2GAPB and RASAL3 shuts off protrusions by inhibiting Ras and PI3K activities in *Dictyostelium* and neutrophils. Cartoons illustrating mechanism of **(A)** opto-C2GAPB or **(B)** opto-RASAL3 recruitment to the cell front with help of SspB-iLID or CRY2-CIBN optogenetic system, respectively. Time-lapse confocal images of vegetative *Dictyostelium* single cells expressing **(C)** mRFPmars-SspB R73Q-C2GAPB or **(E)** tgRFPt-SspB R73Q-Ctrl (control without C2GAPB). C2GAPB or Ctrl is recruited to the front of the migrating cell by applying 488 nm laser near it, as shown by the dashed white box. White arrows denote existing older protrusions whereas pink arrows highlight emerging newer protrusions. Time in min:sec format. Scale bars represent 5 µm. Polar histogram demonstrates higher probability of fresh protrusion formation away from C2GAPB recruitment area (n_c_=11 and n_p_=45) **(D)** whereas for Ctrl, new protrusions form in the same direction as the old ones, near or at the Ctrl recruitment area (n_c_=13 and n_p_=76) **(F)**. **(G, H)** Time-lapse confocal images of vegetative *Dictyostelium* single cells expressing mRFPmars-SspB R73Q-C2GAPB (upper panel) and **(G)** RBD-YFP or **(H)** PHcrac-YFP (both lower panels) after 488 nm laser was switched on globally. RBD or PHcrac status at ‘00:00’ is considered as a control timepoint since C2GAPB has not been recruited yet. Pink arrows denote RBD or PHcrac patches in the cells. Time in min:sec format. Scale bars represent 5 µm. **(I, J)** Time-lapse confocal images of *Dictyostelium* electrofused giant cells expressing mRFPmars-SspB R73Q-C2GAPB (upper panel; red) and **(I)** RBD-YFP or **(J)** PHcrac-YFP (both lower panels; yellow) before and after 488 nm laser was switched on globally. White arrows highlight C2GAPB recruitment in the red channel whereas pink arrows denote RBD or PHcrac waves near the bottom surface of these cells. Time in min:sec format. Scale bars represent 10 µm. **(K)** Time-lapse confocal images of differentiated HL-60 neutrophil expressing CRY2PHR-mCherry-RASAL3 (red; upper panel) and LifeAct-miRFP703 (cyan; lower panel). RASAL3 was recruited to the cell front by applying 488 nm laser near it, as shown by the dashed white box. White arrows denote existing older protrusions whereas pink arrows highlight emerging newer protrusions. Time in min:sec format. Scale bars represent 5 µm. **(L)** Polar histogram demonstrates higher probability of fresh protrusion formation away from RASAL3 recruitment area; n_c_=12 and n_p_=30.

Since C2GAPB had a strong effect on cell behavior, we checked whether it was having this effect by reducing Ras and PI3K activities. While unrecruited cells displayed multiple, small RBD or PHcrac patches (‘00:00’ in Figure 2G and H), once C2GAPB was rapidly recruited it caused a simultaneous reduction in activated Ras and PIP3 levels at the protrusions (‘00:15’-‘2:30’ in Figure 2G and H). Similarly, we observed a strong reduction in activated Ras and PIP3 propagating waves upon C2GAPB recruitment (Figure 2I and J; Videos S5 and S6). Once we switched off the blue light, RBD waves recovered within a minute (Video S5).

We extended our investigation to differentiated HL-60 neutrophils to assess the conservation of RasGAP function on cytoskeletal remodeling. As previously reported, we developed a recruitable RASAL3 by fusing it with CRY2-mCherry, allowing for recruitment to the membrane anchor, CIBN-CAAX (Figure 2B)^4^. Recruiting RASAL3 to the F-actin-rich front, marked by LifeAct, caused mature protrusions to immediately disappear. Simultaneously, a new broad front emerged at the opposite end of the cell, causing it to migrate away (Figure 2K; Video S7). Angular histogram analysis confirmed that new protrusions consistently appeared at ∼130-170 degrees from the RASAL3 recruitment site (Figure 2L).

These observations, both in *Dictyostelium* and human neutrophils, demonstrate that locally recruiting RasGAPs resulted in the immediate shutdown of front signaling, and consequently cellular protrusions, and thereby reversed the existing front-back polarity.

### Inhibitory function of RasGAPs induce polarity and enhance random cell migration

Given the results from Figures 1 and 2, we expected that ectopic global expression of RasGAPs would completely inhibit cellular activity and largely decrease cell migration. Surprisingly, inducing C2GAPB expression resulted in a more polarized phenotype (Figure 3A, also see Figure 1A and B). Moreover, polarized C2GAPB-expressing cells exhibited accelerated movement compared to control cells expressing empty vector, as shown in time series images and color-coded temporal overlay profiles (Figure 3B and C; Video S8). Quantification revealed a 2-fold increase in average cell speed and aspect ratio, a proxy for cell polarity, as compared to uninduced control (Figure 3D and E). Although the C2GAPB-expressing cells changed shape, we did not observe a significant increase in basal cell area (Figure 3F).

**Figure 3.**
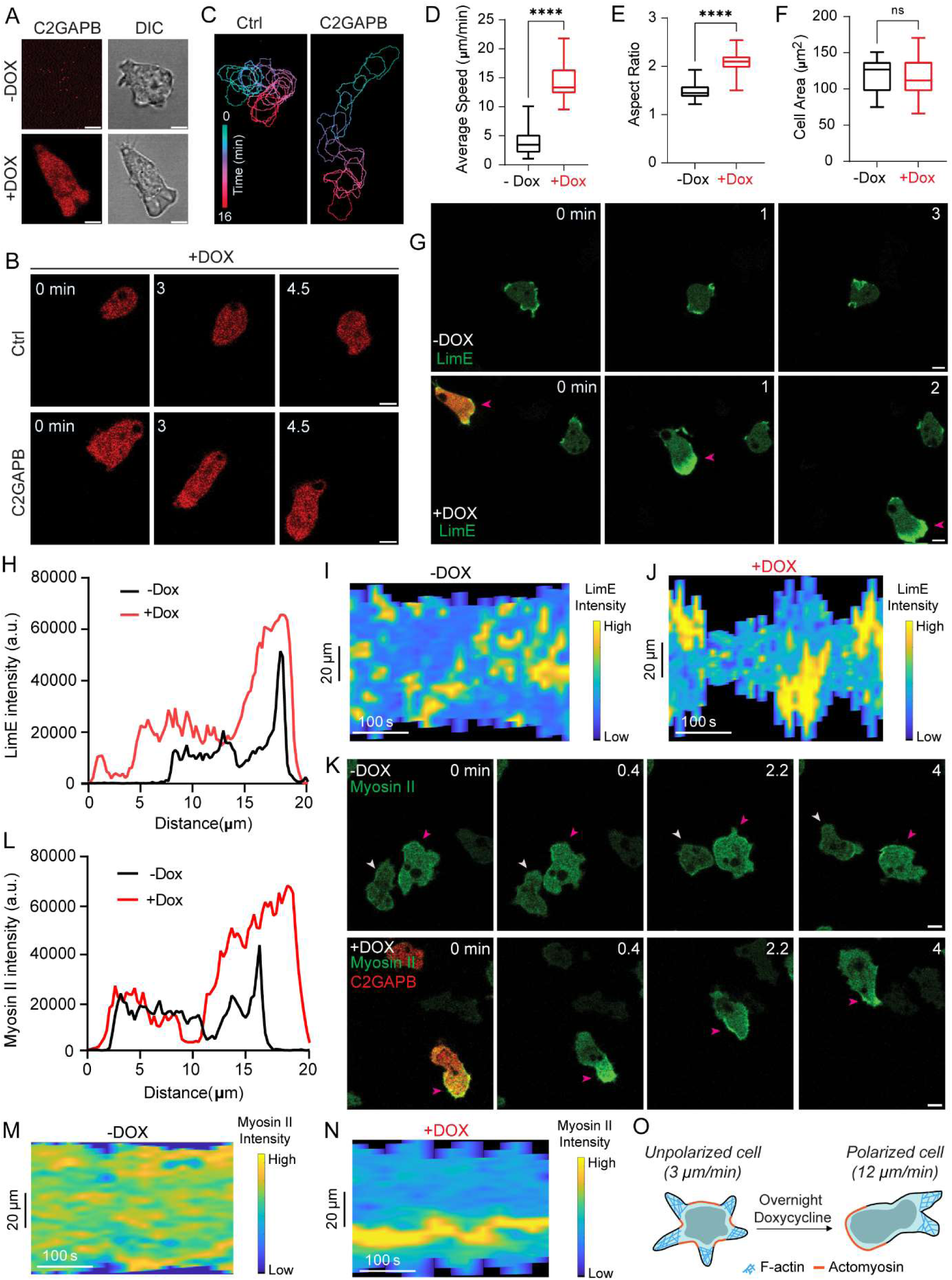
C2GAPB expression polarizes *Dictyostelium* and improves random cell migration. **(A)** Confocal images demonstrating doxycycline-induced mRFPmars-C2GAPB expression (red) polarize vegetative *Dictyostelium* cell (shown in DIC). Scale bars represent 5 µm. **(B)** Time-lapse confocal images of vegetative *Dictyostelium* cells expressing tgRFPt-Ctrl (empty vector control, red; top panel) or mRFPmars-C2GAPB (red; bottom panel) after overnight doxycycline treatment. Time in ‘min’ format. Scale bars represent 5 µm. **(C)** Color-coded (at 1-min interval) outlines of the Ctrl- or C2GAPB-expressing cell shown in (B). Box-and-whisker plots of **(D)** average cell speed, **(E)** aspect ratio, and **(F)** cell area before (black; -DOX) or after (red; +DOX) overnight doxycycline induction of C2GAPB. n_c_=34 from atleast 3 independent experiments; asterisks indicate significant difference, ****P ≤ 0.0001 and ns denotes P>0.05 (Wilcoxon-Mann-Whitney rank sum test). Time in min:sec format. Scale bars represent 5 µm. Time-lapse confocal images of vegetative *Dictyostelium* cells expressing **(G)** GFP-LimE_Δcoil_ (LimE, F-actin biosensor; green) or **(K)** myosin II-GFP (green), before (-DOX) and after (+DOX) overnight doxycycline-induction of C2GAPB (red; not shown here). White or pink arrows denote cells of interest. Time in ‘min’ format. Scale bars represent 5 µm. Representative linescan of **(H)** LimE or **(L)** myosin II intensity of cells in (G) or (K) respectively. Representative kymograph of cortical **(I, J)** LimE or **(M, N)** myosin II intensity in cells in (G) or (K) respectively. A linear color map shows that blue is the lowest LimE or myosin II intensity whereas yellow is the highest. **(O)** Cartoon summarizing the polarizing effects of expressing C2GAPB in *Dictyostelium*. Unpolarized cells with small, transient protrusions become polarized with a distinct F-actin front and a myosin II-labelled back after C2GAPB is expressed.

To further characterize this polarization phenotype induced by C2GAPB expression, we examined the localization pattern of two front-back indicators for cytoskeletal activities, LimE, a biosensor for F-actin, and myosin II. Vegetative, wildtype *Dictyostelium* typically displayed multiple, transient, small F-actin fronts, whereas C2GAPB null cells demonstrated an unpolarized morphology with wide fronts (Figures 3G and S1D). However, in C2GAPB-expressing cells, there was a single, broad, polarized LimE-rich protrusion persistently localized at the front of the migrating cell (Figure 3G). Similarly, C2GAPB expression caused myosin II to localize predominantly at the back of the cells, while a more diffused distribution was generally observed in uninduced cells (Figure 3K). These observations were supported by linescan and kymograph analyses (Figure 3H-J and L-N), demonstrating that after C2GAPB induction, actin polymerization and myosin II were localized to a single front and back while being suppressed elsewhere (Figure 3O). Although the acquisition of polarity is a defining characteristic of differentiation in *Dictyostelium*, C2GAPB-induced polarization did not require development or GPCR signaling. C2GAPB expression induced similar polarization in Gβ null cells, which are typically unpolarized with low motility, and improved their migration speed (Figure S2A-E)^41–43^.

Given the surprising polarizing effect of C2GAPB expression, we checked whether recruiting C2GAPB to the cell membrane, in closer association with Ras, would exaggerate its polarization effects. We noticed that expressing the recruitable C2GAPB protein, fused to SspB, only moderately induced polarity (Figure 4B), as compared to the native C2GAPB (Figures 1 and 3). The SspB tag may be responsible for a weaker C2GAPB function, but this was not explored in our study. However, the global recruitment of C2GAPB triggered a robust and ‘instant polarization’ response within a minute resulting in a 2-fold increase in average migration speed and aspect ratio, but not basal cell area (Figure 4A-E; Videos S9 and S10). However, once the blue laser was switched off, cells reverted to their original morphology within minutes (Figure 4F-H; Video S10). There were several F-actin rich fronts in unrecruited cells (‘00:00’ in Figure 4I). C2GAPB recruitment first reduced LimE signals everywhere on the membrane, then as the cell polarized, a single and persistent actin polymerization site grew at the leading front (Figure 4I and J). Interestingly, we also successfully polarized Gβ null cells with C2GAPB recruitment (Figure S2F and G, Video S11). Taken together, these observations also imply that polarity establishment does not require changes in gene and protein profiles which do not take place within a minute.

**Figure 4.**
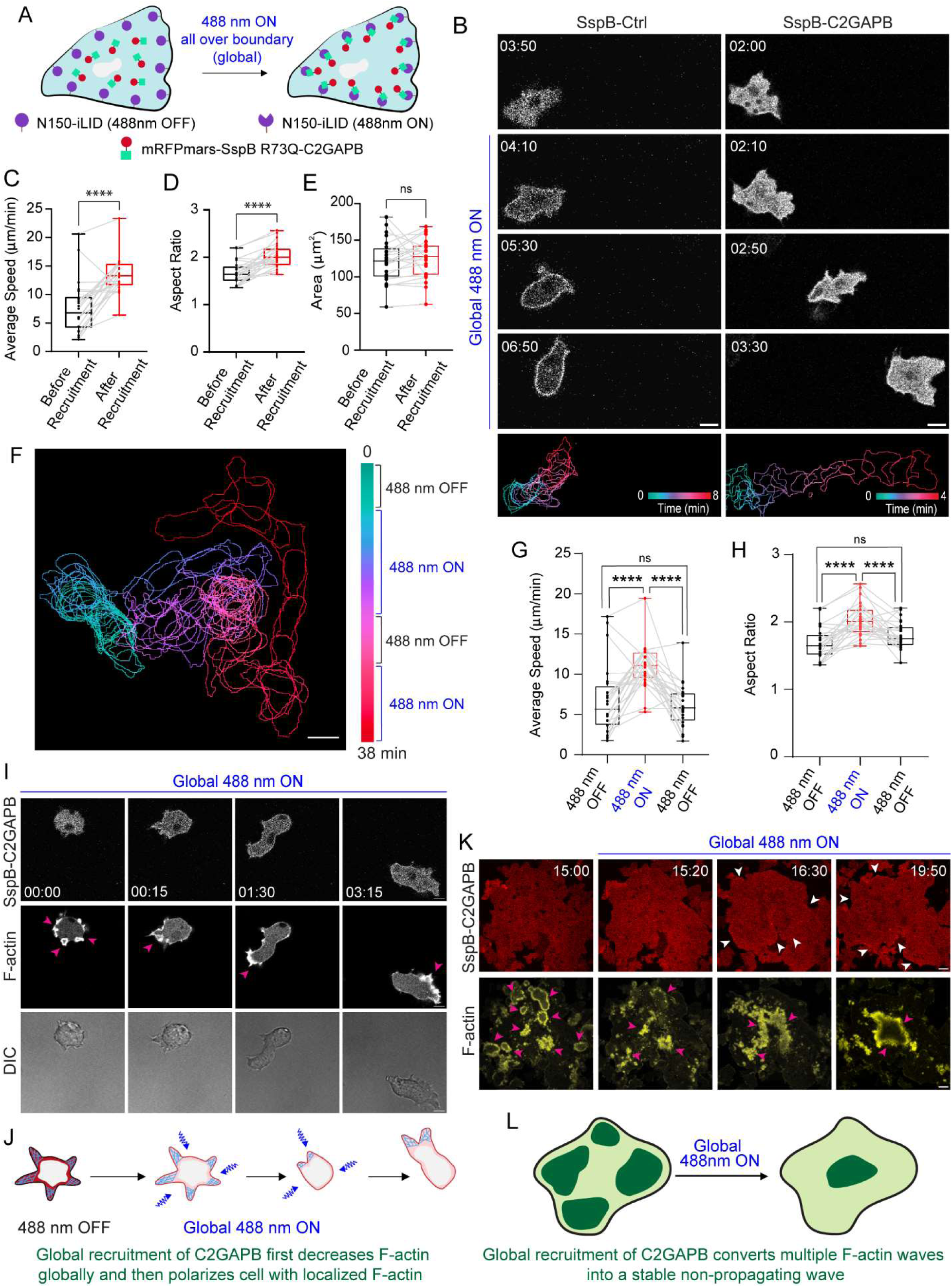
Global C2GAPB recruitment polarizes *Dictyostelium* and enhances cell migration. **(A)** Cartoon illustrating mechanism of opto-C2GAPB global recruitment on *Dictyostelium* cell membrane with help of SspB-iLID optogenetic system. **(B)** Time-lapse confocal images of vegetative *Dictyostelium* cell expressing tgRFPt-SspB R73Q-Ctrl (control without C2GAPB; left panels) or mRFPmars-SspB R73Q-C2GAPB (right panels), before or after 488 nm laser was switched on globally. Bottom panels demonstrate color-coded (at 1-min interval) outlines of the Ctrl- or C2GAPB-recruited cell. Time in min:sec format. **(F)** Color-coded (1-min interval) outlines of a representative cell in presence of intermittent 488 nm light. Scale bars represent 10 µm. Box-and-whisker plots of **(C, G)** average cell speed, **(D, H)** aspect ratio, and **(E)** cell area, before (black) or after (red) C2GAPB global recruitment. For plots in G and H, blue laser was switched on or off multiple times. n_c_=25 from atleast 3 independent experiments; asterisks indicate significant difference, ****P ≤ 0.0001, and ns denotes P>0.05 (Wilcoxon-Mann-Whitney rank sum test). **(I)** Time-lapse confocal images of vegetative *Dictyostelium* single cells co-expressing mRFPmars-SspB R73Q-C2GAPB (upper panel) and LimE-YFP (middle panel) after 488 nm laser was switched on globally. LimE patches are highlighted with pink arrows. ‘00:00’ is considered as control timepoint since C2GAPB has not been recruited yet. DIC channel (bottom panel) shows change in cell polarity with C2GAPB recruitment. Time in min:sec format. Scale bars represent 5 µm. **(J)** Cartoon illustrates phenomenon seen in cell in (I). **(K)** Time-lapse confocal images of electrofused *Dictyostelium* giant cells co-expressing mRFPmars-SspB R73Q-C2GAPB (upper panel; red) and LimE-YFP (lower panel; yellow) before or after 488 nm laser was switched on globally. White arrows highlight C2GAPB recruitment in the red channel whereas pink arrows denote LimE propagating waves near the bottom surface of these cells. Time in min:sec format. Scale bars represent 10 µm. **(L)** Cartoon illustrates phenomenon seen in giant cell in (K).

Next, we validated these C2GAPB-induced effects on LimE wave patterns on the basal surface of electrofused cells. Like single cells, expressing recruitable C2GAPB modestly affected wave patterns (Figure 4K). However, upon recruitment, F-actin waves largely disappeared throughout the cell except for one region on the basal membrane where LimE signal remained strong and did not propagate (Figure 4K and L; Video S12). Thus, our data shows that both signaling, and cytoskeletal waves are largely extinguished but cytoskeletal activity remains persistently confined to one location on the membrane, presumably indicating sustained polarity.

### Localization of recruited RasGAP to the back of the cell led to even stronger polarization

Previously, we showed that recruiting RASAL3 to the front in neutrophils extinguished mature protrusions and made the cells move away (Figure 2K and L). When we continued to extinguish any new protrusions, the cell rounded up completely and did not move (Figure 5A). However, once we stopped extinguishing new protrusions, the cell continued to move away persistently from the last recruitment area (Figure 5A and B; Video S13). Strikingly, RASAL3 simultaneously rearranged itself to the back region, presumably dragging CIBN-CAAX membrane anchor with it. The previously non-polarized, migration-incompetent cell now had a stable back and a sustained front which allowed it to migrate away (Figure 5A-C).

**Figure 5.**
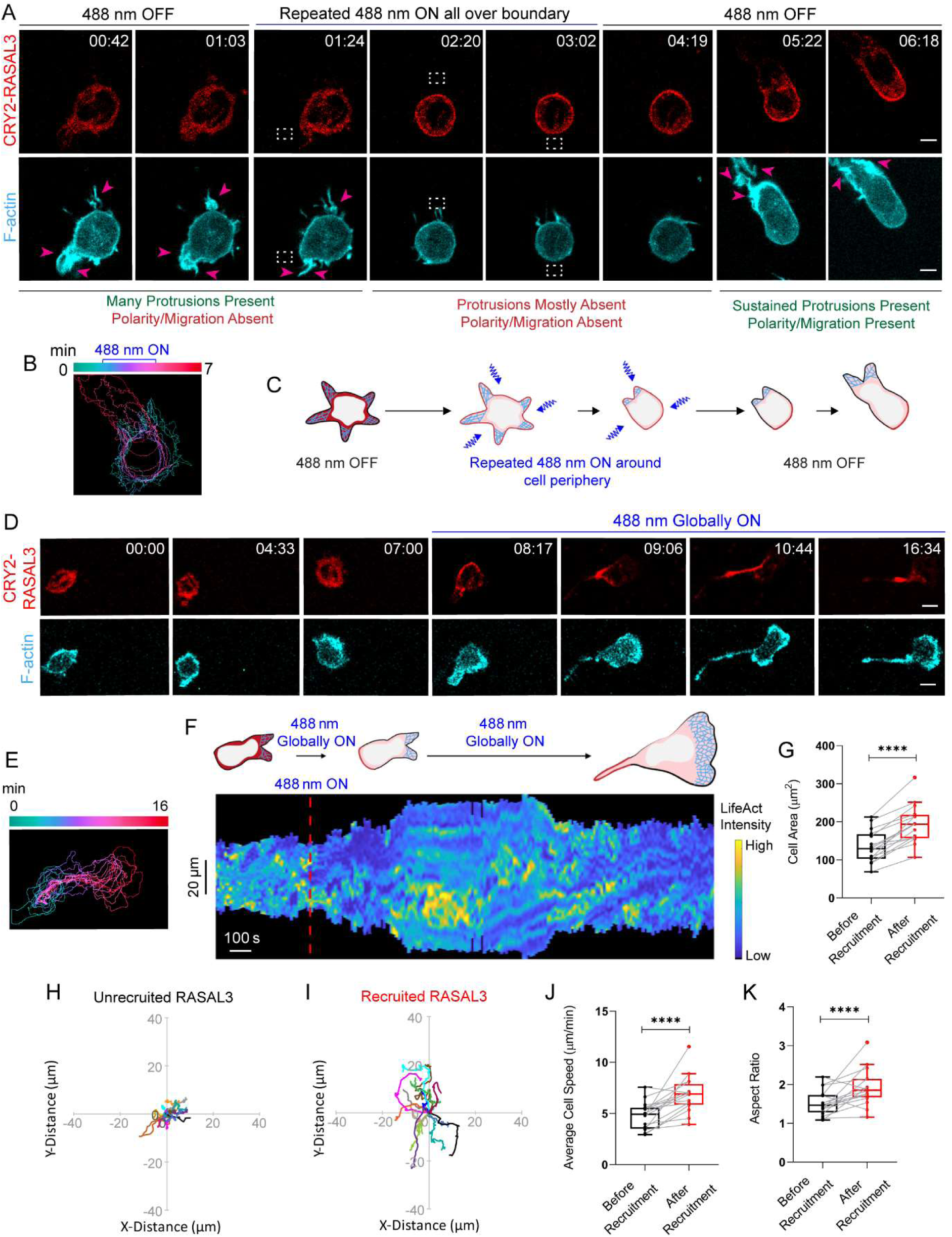
Localization of recruited RASAL3 to the back of the neutrophil led to even stronger polarization. **(A)** Time-lapse confocal images of HL-60 neutrophil expressing CRY2PHR-mCherry-RASAL3 (red; upper panel) and LifeAct-miRFP703 (cyan; lower panel). Unpolarized, non-migratory neutrophil had multiple F-actin-rich protrusions around its perimeter, as shown by pink arrows. RASAL3 recruitment over the entire periphery, after 488 nm light was repeatedly applied all around the cell, caused cell to shrink and LifeAct-containing protrusions soon disappeared. Once blue laser was switched off, RASAL3 self-arranged to the back causing cell to polarize, form LifeAct-rich protrusions, and migrate away. Pink arrows highlight cellular protrusions. Region of blue light illumination is shown by the dashed white box. Time in min:sec format. Scale bars represent 5 µm. **(B)** Color-coded (at 1 min intervals) outlines of the cell shown in (A). **(C)** Cartoon illustrating RASAL3-mediated phenomenon seen in (A). **(D)** Time-lapse confocal images of differentiated HL-60 neutrophil expressing CRY2PHR-mCherry-RASAL3 (red; upper panel) and LifeAct-miRFP703 (cyan; lower panel), before or after 488 nm laser was switched on globally. Time in min:sec format. Scale bars represent 5 µm. **(E)** Color-coded (at 1 min intervals) outlines of the cell shown in (D). **(F)** Representative kymograph of cortical LifeAct intensity in RASAL3-expressing neutrophil before or after 488 nm laser was turned on. A linear color map shows that blue is the lowest LifeAct intensity whereas yellow is the highest. Duration of the kymograph is 24 mins. Cartoon depicts membrane recruitment, F-actin polymerization or cell shape status corresponding to the kymograph. Box-and-whisker plots of **(G)** cell area, **(J)** cell speed, and **(K)** aspect ratio, before (black) or after (red) RASAL3 membrane recruitment. n_c_=16 from atleast 3 independent experiments; asterisks indicate significant difference, ****P ≤ 0.0001 (Wilcoxon-Mann-Whitney rank sum test). Centroid tracks of neutrophils (n_c_=16) showing random motility before **(H)** or after **(I)** RASAL3 membrane recruitment. Each track lasts atleast 5 mins and was reset to same origin.

This spontaneous ‘back’ rearrangement of RASAL3 made it necessary to repeatedly extinguish new fronts to prevent cell movement, since when RASAL3 was globally recruited over the entire cell periphery, it rapidly localized to the rear and polarized the cell. This self-arranged back localization was dependent on cytoskeletal dynamics (Figure S3A and B) and was mediated through the C-terminal tail of RASAL3 (Figure S3C-E). As a result of this strong back localization, neutrophils developed long adhesive uropods at the RASAL3-enriched regions, became highly polarized and migrated (Figure 5D and E; Video S14). Although F-actin levels increased at the front with RASAL3 recruitment, it did not appear to be as high as F-actin changes induced with recruitment of constitutively active Ras or RasGEF shown previously^4^. Across the population, upon RASAL3 back localization, cells moved more extensively with a 1.4-fold increase in average cell speed (Figure 5G-I). Generally, RASAL3 recruitment also induced 1.44- and 1.26-fold increase in cell area and polarity, respectively (Figure 5J and K). A similar polarization was induced upon recruiting the GAP domain fused to its C-terminal tail (Figure S3F-H). No such phenotypic change was observed when the CRY2PHR component, without being fused to RASAL3, was recruited to the cell membrane (Figure S4A-C; Video S15; also Figure S3H-N in^4^). Similarly, when we recruited C2GAPB locally to the back of *Dictyostelium*, the cell moved away persistently (Figure S4D and E). Altogether, these results suggest that reduction in Ras activity can polarize cells, but the localization of RasGAPs to the back can lead to a strong polarization of cells.

### RasGAPs polarize cells by increasing contractility at the back

We were interested to check if the RasGAP-induced protrusions were derived from Arp2/3-mediated actin. We used Arp2/3 inhibitor, CK-666, which completely removed all protrusions in neutrophils (Figure S5A)^4,44^. When we globally recruited RASAL3 and it localized to the back, interestingly, cells made long, thin protrusions which did not display LifeAct biosensor at the tips. This led us to believe that these structures are long blebs which were forming presumably through hydrostatic pressure elsewhere. Rather, the biosensor appeared to be along the lateral edges of these narrow protrusions (Figure S5A and B). Similarly, long blebs were induced upon C2GAPB expression in CK-666-treated *Dictyostelium* cells, but not in control cells where C2GAPB was absent (Figure S5C and D). These observations suggested that the polarizing effects of the RasGAPs were due to increased contraction.

To test whether this contraction was mediated by actomyosin, we induced C2GAPB expression in myosin II heavy chain null (mhcA^-^) mutant of *Dictyostelium*, and also recruited C2GAPB to the membrane. The mhcA^-^ cells, as previously reported, are generally flattened, unpolarized, and multinucleated cells^45–48^. Expression of C2GAPB had little effect on improving this existing phenotype in this cell line (Figure 6A). Furthermore, robust recruitment of C2GAPB that typically polarizes wild type cells, failed to do so in the mhcA^-^ cells (Figure 6B). Similarly, in C2GAPB-recruited polarized cells, blebbistatin (myosin II inhibitor) could diminish polarity although it was not as effective as a complete loss of myosin in the mhcA^-^ cells (Figure S6A)^49^.

**Figure 6.**
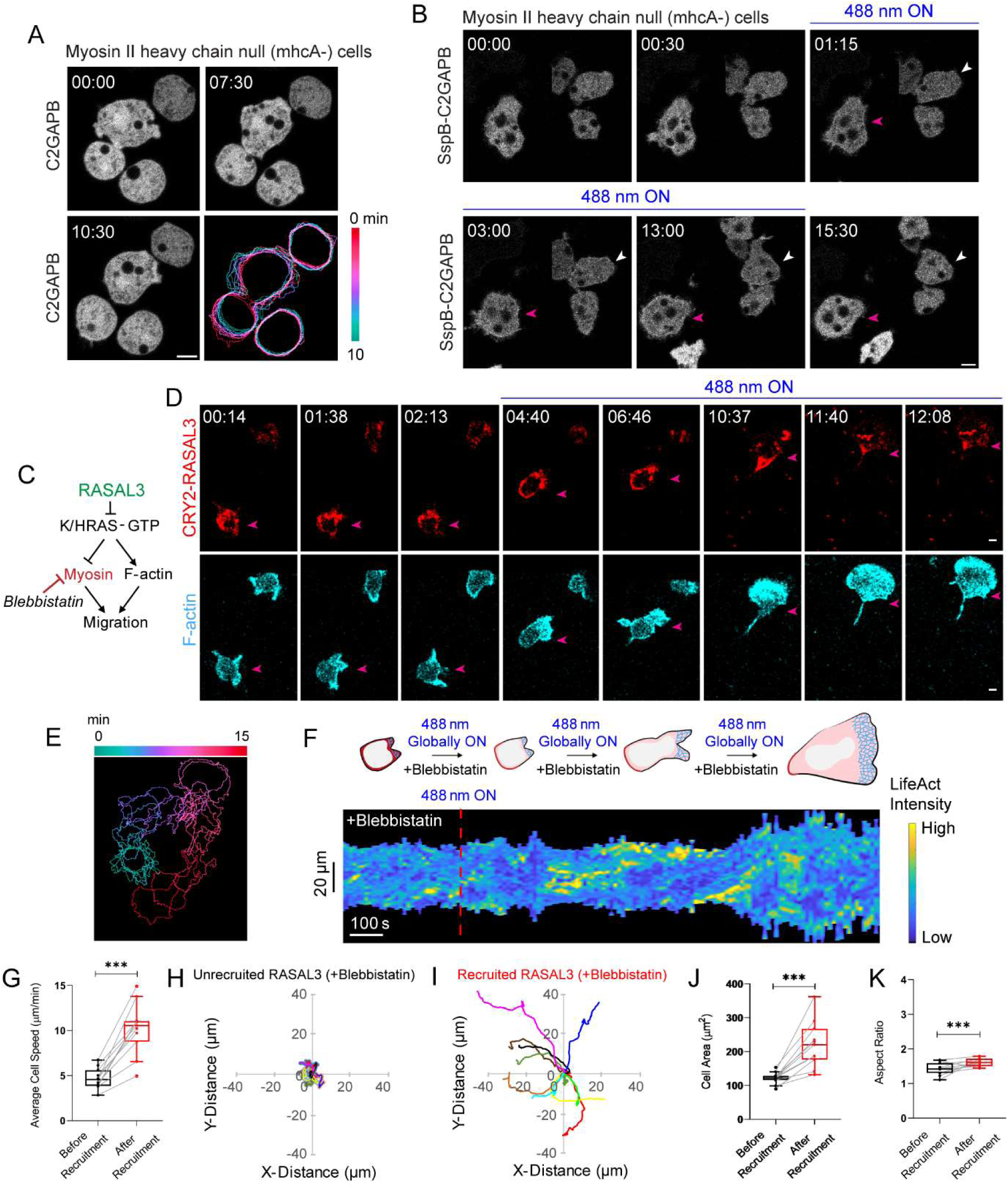
RasGAPs polarize cells by increasing actomyosin contractility at the back. **(A)** Time-lapse confocal images of vegetative myosin II heavy chain null (mhcA^-^) *Dictyostelium* cells expressing C2GAPB after overnight doxycycline treatment. Time in min:sec format. Scale bars represent 5 µm. Color-coded (at 1-min interval) outlines of C2GAPB-expressing cells. **(B)** Time-lapse confocal images of mhcA^-^ *Dictyostelium* cell expressing mRFPmars-SspB R73Q-C2GAPB, before or after 488 nm laser was switched on globally. White and pink arrows denote cells of interest. Time in min:sec format. Scale bars represent 5 µm. **(C)** Strategy for testing effects of myosin inhibitor, blebbistatin, on RASAL3-mediated actomyosin contraction and migration. **(D)** Time-lapse confocal images of blebbistatin-treated HL-60 neutrophil expressing CRY2PHR-mCherry-RASAL3 (red; upper panel) and LifeAct-miRFP703 (cyan; lower panel), before or after 488 nm laser was turned on globally. Pink arrows denote cell of interest. Time in min:sec format. Scale bars represent 5 µm. **(E)** Color-coded (at 1 min intervals) outlines of the cell shown in (D). **(F)** Representative kymograph of cortical LifeAct intensity in blebbistatin-treated RASAL3-expressing neutrophil before or after 488 nm laser was turned on. A linear color map shows that blue is the lowest LifeAct intensity whereas yellow is the highest. Duration of the kymograph is 15 mins. Cartoon depicts membrane recruitment, actin polymerization or cell shape status corresponding to the kymograph. Box-and-whisker plots of **(G)** cell speed, **(J)** cell area, and **(K)** aspect ratio, before (black) or after (red) RASAL3 membrane recruitment. n_c_=12 from atleast 3 independent experiments; asterisks indicate significant difference, ***P ≤ 0.001 (Wilcoxon-Mann-Whitney rank sum test). Centroid tracks of blebbistatin-treated neutrophils (n_c_=12) showing random motility before **(H)** or after **(I)** RASAL3 membrane recruitment. Each track lasts atleast 5 mins and was reset to same origin.

We tested whether myosin-based contractility was needed in neutrophils by treating them with blebbistatin (Figure 6C). Treatment with the myosin II inhibitor resulted in complete stoppage of cell movement and loss of polarity (Figures 6D and S6B)^50–52^. When RASAL3 was globally recruited, it moved to a particular region which became the new back. This local suppression of Ras activity in the region overcame blebbistatin inhibition and caused increased contraction. The polarized cells began to move rapidly, although the blebbistatin-treated cells looked and behaved differently from untreated, RASAL3-recruited cells (Figure 6D; Video S15). With RASAL3 recruitment, compared to untreated cells, blebbistatin treated cells migrated faster with largely diminished uropods and bigger F-actin-driven fronts (Figure 6D and E, G-I; Video S16). Representative kymograph showed increased LifeAct intensity and cell area (Figure 6F). Overall, blebbistatin-treatment improved migration speed and cell area of RASAL3-recruited cells by over 50% and 25%, respectively (Figures 6G-J and 5G-J), whereas aspect ratio was reduced to half of what was seen for untreated, recruited cells (Figures 6K and 5K). We validated our results by treating neutrophils with the ROCK inhibitor, Y27632, which stalled basal motility (Figure S6C and D)^53–55^. RASAL3 recruitment and its back localization mostly overcame the effects of ROCK inhibition; cells polarized and migrated with huge F-actin rich fronts and relatively short uropods (Figure S6E-G; Video S17). Generally, RASAL3 recruitment induced 1.91-, 1.53- and 1.46-fold increase in cell speed, cell area and polarity, respectively (Figure S6H-L). While the inability of C2GAPB to polarize *Dictyostelium* mhcA^-^ cells suggests that myosin is required, it is unclear whether myosin is dispensable for RASAL3 to polarize neutrophils. It is possible that the inhibitors used are not completely effective. Nevertheless, our data suggests that myosin does contribute to the rear contraction since RASAL3-induced uropods are much weaker in the presence of blebbistatin or Y27632^56^.

### RasGAPs polarize cells by localizing actin polymerization at the front

Previously, we observed that back localization of recruited RASAL3 induced F-actin to localize to a stable front which caused neutrophils to polarize and migrate persistently (Figure 5D-K). To confirm that RASAL3 at the back is directly affecting protrusive activity at the front, we pharmacologically targeted front signaling pathways in neutrophils before recruiting RASAL3. First, we inhibited PI3K/PIP3 signaling using pan-PI3K inhibitor, LY294002, which stalled neutrophil polarity and basal motility (Figure 7A and B)^4,57^. Within 5 minutes of applying blue light globally, RASAL3 moved to the back region of LY294002-treated cells and polarized them to move by generating broad F-actin-rich lamellipodium but without an appreciable uropod (Figure 7B and C, F and G; Video S18). The kymograph shows that the LifeAct level increased considerably along with an increase in cell area (Figure 7D). Across the entire population, RASAL3 recruitment caused 1.75-, 1.49, or 1.43-fold improvement in average cell speed, basal cell area or aspect ratio (Figure 7E, H and I). We observed a similar recovery in polarity and random motility in presence of a PI3Kγ inhibitor, AS605240, with RASAL3 recruitment (Figure S7; Video S19)^4,58,59^.

**Figure 7.**
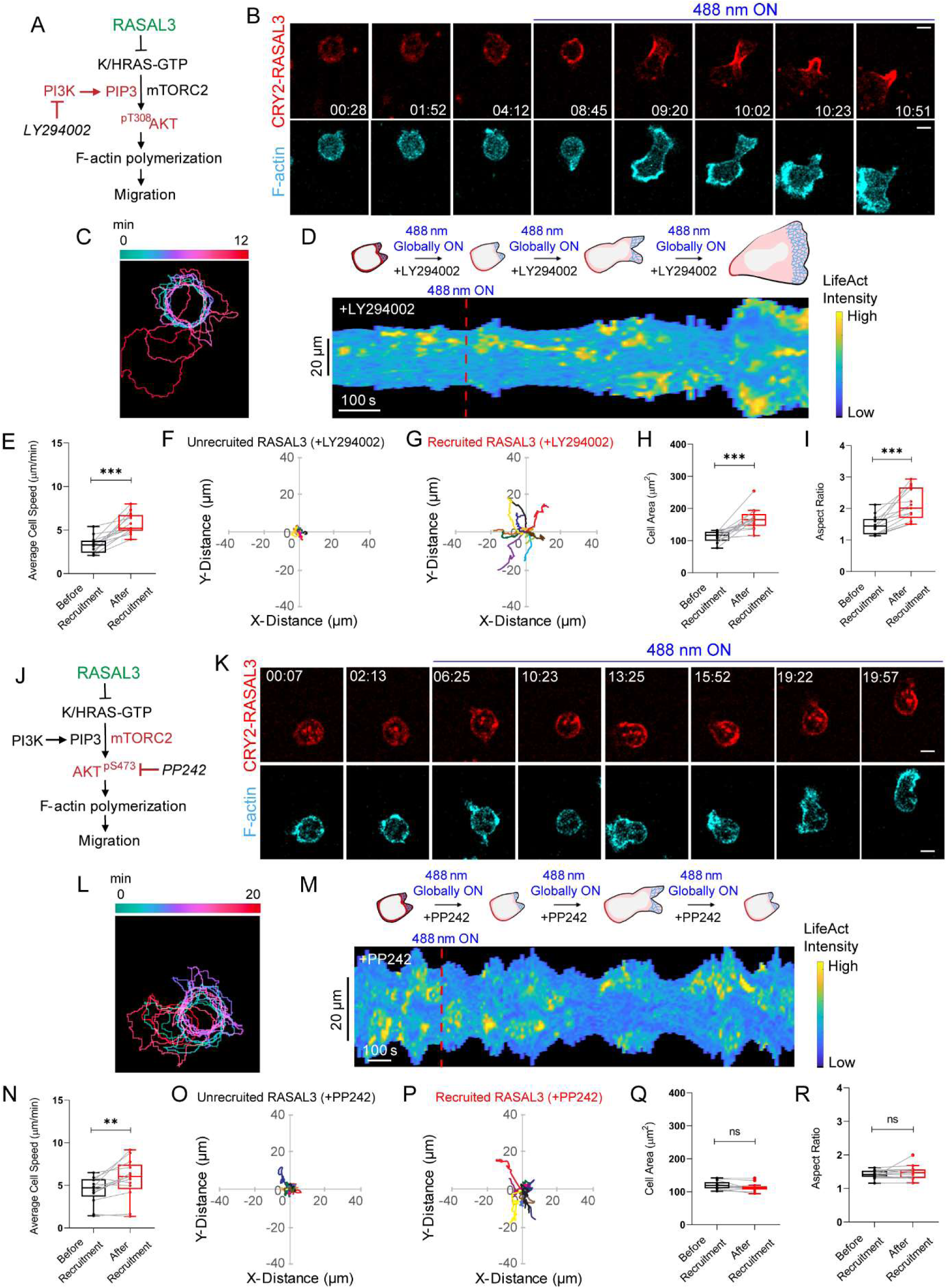
RASAL3 polarizes cells by localizing actin polymerization at the front. Strategy for testing effects of **(A)** pan-PI3K inhibitor, LY294002, or **(J)** mTOR inhibitor, PP242 on RASAL3-directed actin polymerization and motility. Time-lapse confocal images of **(B)** LY294002- or **(K)** PP242-treated HL-60 neutrophil expressing CRY2PHR-mCherry-RASAL3 (red; upper panel) and LifeAct-miRFP703 (cyan; lower panel), before or after 488 nm laser was turned on globally. Time in min:sec format. Scale bars represent 5 µm. **(C, L)** Color-coded (at 1 min intervals) outlines of the cells shown in (B) and (K), respectively. Representative kymographs of cortical LifeAct intensity in **(D)** LY294002- or **(M)** PP242-treated RASAL3-expressing neutrophil before or after 488 nm laser was turned on. A linear color map shows that blue is the lowest LifeAct intensity whereas yellow is the highest. Duration of the kymographs are 12 mins and 20 mins, respectively. Cartoons depict membrane recruitment, actin polymerization or cell shape status corresponding to the kymographs. Box-and-whisker plots of **(E or N)** cell speed, **(H or Q)** cell area, and **(I or R)** aspect ratio, before (black) and after (red) RASAL3 recruitment in LY294002- or PP242-treated cells. n_c_=13 from atleast 3 independent experiments; asterisks indicate significant difference, ***P ≤ 0.001, **P ≤ 0.01, and ns denotes P>0.05 (Wilcoxon-Mann-Whitney rank sum test). Centroid tracks of LY294002- or PP242-treated neutrophils (n_c_=13) showing random motility before **(F or O)** or after **(G or P)** RASAL3 recruitment. Each track lasts atleast 5 mins and was reset to same origin.

Next, we inhibited mTORC2 signaling in neutrophils using the established mTOR inhibitor, PP242 (Figure 7J)^60^. Inhibitor treatment caused cells to round up, which is due to mTORC2 complex inhibition and not mTORC1 (Figure 7K)^4,61^. Once RASAL3 recruitment was initiated after turning on the blue laser, cells displayed weak, intermittent polarity (Figure 7K and L). Time-lapse imaging and migration tracks showed that recruited RASAL3, which goes to the back, moved PP242-treated cells slightly, but not persistently (Figure 7K, O and P; Video S20). As a result, although we noticed a 1.3-fold improvement in average cell speed, we did not see any significant change in cell area or polarity (Figure 7M and N, Q and R).

### Increasing RasGAP in simulations of the Signal Transduction Excitable Network lead to reduced Ras activity but cells can still polarize

To determine the role of RasGAP on the level of Ras on the signaling network, we turned to a model of the excitable behavior that has been used to recreate the observed wave behavior in cells^37,62,63^. As shown in Figure 8A, the model incorporates complementary inhibition between Ras and PIP2 which acts as a positive feedback loop^11,64^. There is a competing negative feedback loop involving Ras and PKB^37,64^. These three elements form the core of the Signal Transduction Excitable Network (STEN). Additionally, we incorporate two further feedbacks to capture the polarizing behavior that is observed over time^62,65^. Note that here we have implemented one of the polarity loops as a positive loop emanating from back events in STEN, such as PIP2, rather than as negative feedback from F-actin. Simulations based on this model display characteristic excitable behavior, including propagating waves that annihilate when they come together (Figure 8B; Video S21). We considered the effect of increasing/decreasing RasGAP activity. Whereas simulations in which RasGAP was reduced showed increased Ras activity and resulted in large, fast wave patterns compared to WT levels of RasGAP, those with increased RasGAP showed small, slow waves (Figure 8B; Video S21). These observations were confirmed by summing all Ras activity throughout the simulations (Figure 8C). The two-dimensional simulations described above can model the basal surface of large, electrofused cells. We also carried out simulations of a single cell, in which activity is measured along the perimeter (Figure 8D). Since polarity is more obvious in these cells, the strength of the feedback loops was increased. In these simulations, we saw great increases in activity before changes in the RasGAP level. Halfway through the simulation, this was increased. This led to an overall decrease in Ras activity. Thereafter, single streaks representing one or two stationary waves were observed (Figure 8D). These observations are similar to the experiment in Figure 3M and 3N.

**Figure 8.**
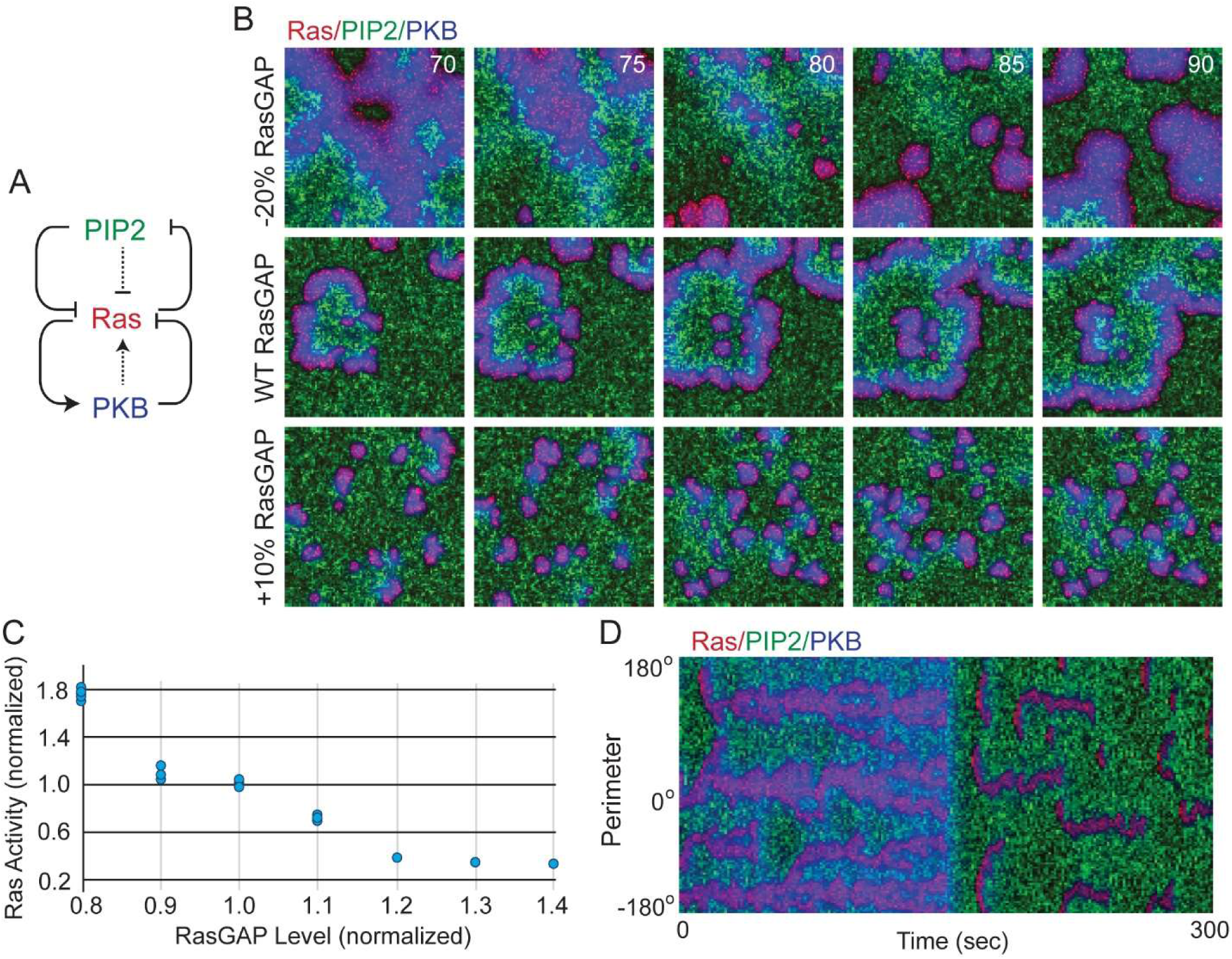
Simulations of the Signal Transduction Excitable Network show increasing RasGAP leads to reduced Ras activity but cells still polarize. **(A)** An excitable system incorporating positive and negative feedback loops between Ras, PIP2, and PKBA was simulated. Also included are two extra loops (dotted lines) described the effect of polarization. **(B)** The EN was simulated under varying levels of RasGAP (Methods). Assuming a wild-type level of RasGAP, firings of the excitable network were seen and led to propagating waves of high Ras activity (red) moving at speeds of ∼10 µm/min. The trailing edge of these wave fronts was marked by high PKB (blue) activity, and by an absence of PIP2 (green). These waves showed characteristic excitable medium behavior, including annihilation of fronts following a collision, wave splitting and spiral activity. In simulations in which the level of RasGAP was lowered by 20%, the size and speed of the wave increased by ∼50% and ∼100%, respectively. A 10% increase in the levels of RasGAP had the opposite effect. Firings were still seen, but they broke up frequently, leading to ∼75% smaller and ∼40% slower waves. The simulation denotes a square where each side is 40 µm and the time is in seconds. **(C)** To quantify the effect of RasGAP activity in the simulations (n=5 for each condition), we computed the total level of Ras over the simulations, while varying RasGAP from 80-140% of wild type levels. Shown is the mean level of activity over the whole simulation. Note that after about a 30% increase in RasGAP, no firings were observed, and hence the level of Ras plateaus. **(D)** We also carried out simulations in smaller cells represented by a one-dimensional environment (representative of the perimeter of a cell). The strength of the RasGAP contribution was increased 50% halfway during the simulation.

## Discussion

Using optogenetic activation techniques on RasGAPs, C2GAPB and RASAL3, expressed in *Dictyostelium* and neutrophils, respectively, our study yielded significant insights. Notably, locally dampening Ras activity could extinguish cellular protrusions and impede migration. However, contrary to expectations, a global reduction in Ras activity heightened both polarity and motility. Furthermore, in *Dictyostelium,* targeting C2GAPB to the cell back amplified migration and polarity, whereas in neutrophils, spontaneous movement of RASAL3 to the back generated uropods and localized F-actin protrusions at the front. Thus, surprisingly, polarity can be achieved by suppressing Ras activity at the rear of the cell. Although Ras and its downstream pathways are typically associated with longer-term growth control, this investigation establishes the immediate and spatially defined function of Ras activity at the cell cortex in governing cell polarity and motility.

How can Ras inhibition improve polarity? We propose that a global decrease in Ras activity raises the threshold for initiating protrusive activity but this is not enough to block all protrusions. Our observation on the effects on RBD or PHcrac wave activity with RasGAP strongly suggests that we are raising the activation threshold of our system. The increased threshold causes typically broad, propagating cortical waves to break up into smaller, short-lived waves in electrofused cells. Analogously, in single cells, RasGAPs suppress multiple, co-existing protrusions all around the periphery and confine them to a single protrusion. This singular protrusion appeared to be further enhanced by some positive feedback. Obviously, if all protrusions were shut down, that would block migration; however, we were unable to achieve this in our study presumably because our RasGAPs were not strong enough to raise the activation threshold enough. In our model, Ras plays a central role in regulating two strong opposing feedback loops consisting of molecules, such as myosin and PI(4,5)P2 that typically form the back, and PIP3 and F-actin that generate the front. This model faithfully recapitulates the results presented here as well as predicting the previously observed increased cell spreading and front activity upon knocking out RasGAPs^8,11,18,22,23^. It also predicts a presumed ability of a further reduction of Ras activity to eliminate cell movement.

Previous studies have suggested that increases in actomyosin contraction at the back, occurring spontaneously or locally triggered by optogenetic RhoA or RGS4, can initiate polarity and cause cells to move away^66–74^. Our study demonstrates that spatiotemporal regulation of Ras signaling pathways not only controls F-actin protrusive activity at the front, but also directly coordinates contraction at the back. Even when Arp2/3 was inhibited, the ability of back suppression to polarize a cell could be observed. Since F-actin was absent, instead of making normal protrusions, suppressing Ras activity at the back led to formation of long bleb-like structures.

In addition to increasing contractility at the back, Ras inhibition at the back appeared to trigger increased F-actin polymerization at the front. This was evidenced by the fact that while inhibitors of signaling downstream of Ras suppressed protrusions and stopped cells from moving, the inhibition could be partially overcome by recruiting RasGAP to the back. These RasGAP-mediated long-range effects on the cell front were mediated primarily through mTORC2, rather than PI3K/PIP3. Traditionally, these possibilities have been difficult to distinguish since conventional chemotactic gradient studies lack sufficient spatiotemporal resolution to determine chronology of front and back formation during symmetry breaking^71,72,74,75^. However, our results revealed that suppression at the back can trigger polarization, which may simultaneously activate the front.

Our study sheds light on the complexity of Ras function in cells. Previous reports have shown that expression of oncogenic Ras leads to excessive protrusions in multiple different cell types^4,20,61^. The extraneous protrusions impair cell movement. Our recent findings clearly show that local activation or inhibition of Ras at the cell perimeter can quickly elicit or extinguish protrusions, indicating that Ras activity is necessary and sufficient^4^. It was, therefore, surprising that global inhibition of Ras activity quite effectively polarized both amoebae and neutrophils. Polarization occurred even though in some of the amoebae experiments Ras activity was reduced to undetectable levels in single cells. Small RBD puncta were still visible in giant, electrofused cells indicating Ras was not completely suppressed with C2GAPB recruitment. Altogether, the studies suggest that, for effective motility, there is an optimal level of Ras activity which is very low. Cells move most effectively when maintaining persistent Ras activation at a narrow location and strongly inhibit on the remainder of the perimeter. Consistently, our modeling showed that polarity was best achieved when Ras activity was strongly reduced. Of course, in the simulation a complete loss of Ras activity could stop movement; we were unable to achieve this experimentally.

Such findings have important implications for cancer treatment, where targeting Ras or constitutively active Ras mutations may not always be beneficial. Caution must be exercised to avoid, while attempting to abrogate cell proliferation, forcing cells into a more polarized migratory state. The increased migration could, for example, induce cells to exit epithelium and metastasize. Consistently, it is well-known that metastasizing cells do not readily divide^76–79^. A deeper understanding of the different roles that Ras activities play in cell migration versus cell growth is essential for developing effective therapeutic strategies.

## Acknowledgements

We thank all members of the Peter Devreotes, Pablo Iglesias, and Douglas Robinson laboratories (Schools of Medicine and Engineering, JHU) for helpful discussions and providing resources. We acknowledge Orion Weiner (UCSF) for providing HL-60 cell line. We thank Sean Collins (UC Davis), Marc Edwards (Amherst College), and Yuchuan Miao (Harvard Medical School) for providing plasmids. We thank Stephen Gould (School of Medicine, JHU) for help with instrumentation. We thank Xiaoling Zhang (Ross Research Flow Cytometry Core, JHU) for helping with cell sorting. We appreciate DictyBase and Addgene for plasmids. This work was supported by NIH grant R35 GM118177 (to P.N.D.), DARPA HR0011-16-C-0139 (to P.A.I. and P.N.D.), AFOSR MURI FA95501610052 (to P.N.D.), as well as NIH grant S10OD016374 (to S. Kuo of the JHU Microscope Facility).

## Author Contributions

DSP, YL, and PND conceived and designed the project; YL and DSP engineered constructs/stable cell lines, designed and executed experiments, and performed majority of data analyses with input from other authors; PB and PAI devised and conducted computational simulations; TB and GQ executed kymograph and some MATLAB/Python analyses; YD performed uptake assays; JB made some constructs; DSP, PND, and YL extensively revised ChatGPT-generated initial drafts, which were developed from bullet points supplied by authors, and wrote final version of the manuscript with help from other authors; DSP and PND supervised the study.

## Competing Interests

The authors declare no competing interests.

## Materials and Methods

### Reagents and inhibitors

200 µg/mL fibronectin stock (Sigma-Aldrich; F4759-2MG) was prepared in sterile water, followed by dilution in PBS. 20 mM AS605240 (Sigma-Aldrich; #A0233), 20 mM PP242 (EMD Millipore; #475988), 50 mM LY294002 (Thermo Fisher; #PHZ1144), 50 mM CK-666 (EMD Millipore; #182515), 50 mM blebbistatin (Peprotech; #8567182), 10 mM latrunculin B (Sigma-Aldrich; #428020), or 5 mM Y27632 (Sigma-Aldrich; #688001) stock solution was made in DMSO (Sigma Aldrich; #D2650). Jasplakinolide (Sigma-Aldrich; #420127) was available as a ready-made 1 mM stock. Hygromycin B (Thermo Fisher Scientific; #10687010) or G418 sulphate (Thermo Fisher Scientific; #10131035) was purchased as 50 mg/mL stock solution whereas blasticidine S (Sigma-Aldrich; #15205) or puromycin (Sigma-Aldrich; #P8833) was dissolved in sterile water to make stock solutions of 10 mg/mL or 2.5 mg/mL, respectively. Doxycycline hyclate (Sigma; #D9891-1G) was dissolved in sterile water to make a stock of 5 mg/mL. 50 mg/mL TRITC-dextran (Sigma-Aldrich; #T1162) was made in sterile water. All stock solutions were aliquoted and stored at -20℃. According to experimental requirements, further dilutions were made in development buffer (DB), PBS, or growth medium before adding to cells.

### Plasmid construction

All DNA oligonucleotides were purchased from Sigma-Aldrich. *Dictyostelium* C2GAPB (RasGAP2, RG2) gene was previously cloned in KF2 expression plasmid^11^. Using this construct, we subcloned C2GAPB into doxycycline-inducible pDM335 plasmid (DictyBase #523) using BglII/SpeI restriction digestion to generate mRFPmars-C2GAPB/pDM335 and GFP-C2GAPB/pDM335 constructs. SspB R73Q ORF was amplified from tgRFPt-SspB R73Q plasmid (Addgene #60416) and then subcloned into C2GAPB/pDM335 at the BglII site to generate the optogenetically-recruitable C2GAPB construct, mRFPmars-SspB R73Q-C2GAPB^80^. Similarly, tgRFPt-SspB R73Q ORF was introduced into pDM335 to generate tgRFPt-SspB R73Q-Ctrl construct. The construct for the membrane anchor, N150-Venus-iLID/pDM358, was made previously^4^. This construct was used to subclone PHcrac-YFP, LimE_Δcoil_-YFP, or RBD-YFP ORF to generate dual expressing N150-Venus-iLID/PHcrac-YFP, N150-Venus-iLID/LimE-YFP, or N150-Venus-iLID/RBD-YFP construct, respectively. The shuttle vector, pDM344 (DictyBase #551), was used for this purpose. Constructs for GFP-RBD and GFP-LimE_Δcoil_ were procured from R. Firtel lab (UCSD) and G. Marriott lab (University of Wisconsin-Madison), respectively, whereas myosin-GFP/pDRH was obtained from D. Robinson lab (School of Medicine, JHU)^5,81,82^.

Mammalian constructs, CRY2PHR-mCherry-RASAL3/pPB, CIBN-CAAX/pLJM1, and LifeAct-miRFP703/pLJM1 were generated previously^4^. DNA sequences (2064-3036 or 1276-3036 bases) encoding the last 322 or 585 amino acids of RASAL3 were PCR amplified and cloned into BspEI/NotI sites of the PiggyBac™ transposon plasmid to generate the CRY2PHR-mCherry-RASAL3_689-1011_/pPB or CRY2PHR-mCherry-RASAL3_426-1011_/pPB construct, respectively (PiggyBac™ transposon system was gifted by S Collins lab, UC Davis)^83,84^.

All constructs were verified by diagnostic restriction digestion and sequenced at the JHMI Synthesis and Sequencing Facility.

### Cell culture

Wild-type *Dictyostelium discoideum* cells of the AX2 strain, obtained from the R. Kay lab (MRC Laboratory of Molecular Biology, UK), were used in this study. The Gβ-null (Gβ^-^) cells were created in our lab previously^41^. Myosin heavy chain-null strain (mhcA^-^) was obtained from D. Robinson laboratory (School of Medicine, JHU)^47^. C2GAPB-null strain (C2GAPB^-^) was generated in our lab^11^. All cell lines were cultured axenically in HL5 medium supplemented with 300 μg/mL streptomycin (GoldBio; #S-150-100) at 22°C. Growth-stage cells were used for all imaging experiments and all strains were used within 2 months of thawing from frozen stocks^85^.

Female human neutrophil-like HL-60 cell line was obtained from Orion Weiner lab, UCSF, and subcultured in RPMI 1640 medium (Gibco #22400-089) supplemented with 15% heat-inactivated fetal bovine serum (FBS; Thermo Fisher Scientific #16140071)^4,86^. To obtain migration-competent cells for experimentation, wildtype or stable cell lines were differentiated in presence of 1.3% DMSO (Sigma #D2650) over 5-7 days^4,86^. Upon differentiation, these cells can be used as an effective model to study human neutrophils^87^. Stable cell lines were maintained in selection antibiotics which were removed during differentiation and experimentation. Cells were grown in humidified conditions at 5% CO_2_ and 37^0^C. Experiments were performed with cells at low passage.

### Stable cell line generation

0.5-1×10^7^ wildtype cells were harvested, washed twice, and finally resuspended in 100 µL chilled H-50 buffer (20 mM HEPES, 1 mM MgSO_4_, 50 mM KCl, 5 mM NaHCO_3_, 10 mM NaCl, and 1 mM NaH_2_PO_4_; pH 7.0). Next, 2 µg total DNA was added to the cell suspension which was subsequently transferred to an ice-cold 0.1-cm electroporation cuvette (BioRad; #1652089), and electroporated at 0.85 kV/25 µF twice with 5 secs between pulses. The cuvette was then incubated on ice for 10 min and cells were transferred to a 10-cm Petri dish (Greiner Bio-One #664160) containing 10 mL HL5 medium supplemented with heat-killed *Klebsiella aerogenes* (lab stock). Next day, 50 μg/mL hygromycin and/or 20 μg/mL G418 sulfate was added, and cells were selected over a period of 3-4 weeks^4,88^.

HL-60 cell line stably co-expressing optically recruitable RASAL3 and LifeAct-miRFP703 was generated previously using a combination of lentiviral- and transposon integration-based approaches^4,88^. In this study, we stably co-expressed recruitable RASAL3_689-1011_ or RASAL3_426-1011_ along with LifeAct-miRFP703 in HL-60 cells using the same protocol.

### Preparation of electrofused giant *Dictyostelium*

Growth-phase cells were harvested, washed, and resuspended in Soerensen buffer (SB; 15 mM KH_2_PO_4_ and 2 mM Na_2_HPO_4_, pH 6.0) at a density of 1.5×10^7^ cells/mL. 7–10 mL cell suspension was put into a 50-mL conical tube and rolled gently for 30 min to promote visible cell cluster formation. 800 μL of ‘rolled cells’ were gently transferred to a 4 mm-gap electroporation cuvette (BioRad; #1652088). Electroporation was then performed using the following settings: 1,000 V, 3 μF once, and 1,000 V, 1 μF thrice, with 1–2 sec between pulses. Next, 50 μL cell suspension was transferred to the center of a well in an 8-well chamber (Lab-Tek, #155409 PK) and left for 5 min. 450 μL SB supplemented with 2 mM CaCl_2_ and 2 mM MgCl_2_ was added to the well and cells were resuspended gently. Medium along with unadhered cells was removed and 450 μL fresh supplemented SB was gently added to the well. Cells were allowed to recover for 1 hr before imaging^35,36^.

### Microscopy

Vegetative *Dictyostelium* in HL5 medium were placed in an 8-well coverslip chamber and allowed to adhere for 40 min. Differentiated HL-60 cells, pre-treated with heat-killed *Klebsiella aerogenes*, were allowed to adhere to fibronectin-coated 8-well coverslip chamber for 40 mins^4,88^. Next, non-adherent cells were washed off, 300-450 μL DB or fresh RPMI 1640 medium was added to the attached *Dictyostelium* or differentiated HL-60 cells, respectively, and used for imaging. To induce C2GAPB expression in *Dictyostelium*, doxycycline (50 µg/mL) was added 8 hr prior to imaging time. For macropinocytosis assay, cells were incubated with TRITC-dextran for 4 mins, washed thrice with DB, and imaged subsequently^33,89^. All time-lapse imaging in *Dictyostelium* or HL-60 cells was acquired with 0.3-0.5% or 1.7-6% laser intensity, respectively, using the following microscopes: (1) Zeiss LSM780-FCS single-point, laser scanning confocal microscope with 780-Quasar; 34-channel spectral, high-sensitivity gallium arsenide phosphide detectors supported with ZEN Black software, and (2) Zeiss LSM800 GaAsP single-point laser scanning confocal microscope with wide-field camera supported with ZEN Blue software. All images of *Dictyostelium* or HL-60 cells were acquired with 63X/1.40 PlanApo oil or 40X/1.30 PlanNeofluar oil DIC objective, respectively, along with digital zoom. In single *Dictyostelium* or HL-60 cells, confocal imaging was performed at a middle plane of the cells, whereas in giant *Dictyostelium* cells, laser was focused near the bottom surface of cells to visualize cortical wave propagation^4,35,88,90^. For inhibitor experiments, differentiated HL-60 cells were treated with 20 µM AS605240, 20 µM PP242, 50 µM LY294002, 50 µM CK-666, 50 µM blebbistatin, or 10 µM Y27632 for atleast 10 mins before imaging. For pre-treatment with JLY cocktail, HL-60 cells were incubated with 10 μM Y27632 for 10 min. These cells were then treated with 8 μM jasplakinolide and 5 μM latrunculin B without changing the final concentration of Y27632^91,92^. In *Dictyostelium*, 50 µM blebbistatin or CK-666 was added during imaging.

Optogenetic experiments with vegetative *Dictyostelium* or differentiated HL-60 cells were done in absence of any chemoattractant. Throughout image acquisition, solid state laser (561 nm excitation and 579-632 nm emission) was used for visualizing proteins or recruitable effectors fused to mCherry, mRFPmars, or tgRFPt tag whereas a diode laser (633 nm excitation and 659-709 nm emission) was used to capture miRFP703 expression. Images were acquired for 5-10 mins, after which 450/488 nm excitation laser was switched on globally to activate recruitment. Image acquisition and photoactivation was done at ∼7 sec intervals. Using the T-PMT associated with the red channel, we acquired DIC images. We took advantage of the interactive photobleaching module on Zeiss LSM800 to perform local recruitment experiments. A small region of interest was placed in front or back of migrating cells, and bleached with 488 nm laser (laser power of ∼0.5% or 7% for *Dictyostelium* or HL-60 cells, respectively) in multiple iteration. Time interval of photoactivation and image acquisition was ∼10 secs. For imaging HL-60 cells, both microscopes were equipped with a temperature-controlled chamber set at 5% CO_2_ and 37^0^C^4,35,88,90^.

### SDS-PAGE and Western blotting

Wildtype or mRFPmars-C2GAPB-expressing *Dictyostelium* was starved and developed, and 6×10^6^ cells were collected every hour, upto 8 hours, for western blot analysis^88,93^. Each sample was resuspended in pre-chilled 1× RIPA buffer (supplemented with protease inhibitor cocktail; Thermo Scientific, # 89900) and lysed on ice for 20 mins. Next, 3× Laemmili sample buffer (lab stock) was added and samples were incubated at RT for 5 mins. Finally, sample equivalent to 1.5×10^6^ cells were loaded into pre-cast 4-15% polyacrylamide gel and was run at 120 V for 1 hour. Immunoblotting was performed as per standardized lab protocol^4,88^. cAR1 expression (∼44 kDa) was detected by incubating PVDF membrane (Bio-Rad; 162-0262) with rabbit anti-cAR1 antibody (1:1000 dilution; generated in our lab^94^) overnight at 4^0^C, followed by goat anti-rabbit IRDye 680RD-cojugated secondary antibody (1:10,000 dilution; Li-Cor; #925-68071) for an hour in the dark. We used Odyssey CLx imaging system (Li-Cor) to detect the near-infrared signal from the blot.

### Image analysis

All images were analyzed with ImageJ (NIH) and MATLAB 2019b (MathWorks, Natick, MA) software^95^. We utilized GraphPad Prism 8 (GraphPad software, CA, USA) and Microsoft Excel (Microsoft, WA, USA) for plotting our results^4,35,88^.

To get the ratio of wave area to cell area in Figure 1E and H, the image was first binarized using ImageJ, by adjusting the threshold to cover all pixels of the wave or cell. The range was not reset and the ‘Calculate threshold for each image’ option was unchecked. Subsequently, using the “Analyze Particle” function, a size-based thresholding was applied (to exclude non-cell particles) and cell masks were generated. Next, the ‘Fill holes’, ‘Erode’ and ‘Dilate’ options were applied, sequentially and judiciously, to obtain the proper binarized mask for waves or cells^4,35^. Next, the ratio of wave area to cell area for each frame was obtained and plotted with time. Duration of waves for Figure 1F and I was obtained by counting the number of frames from when a wave starts to when it ends, and then multiplying it with the time interval.

Cell outline overlays (Figures 3C, 4B, 4F, 5B, 5E, 6A, 6E, 7C, 7L, S2C, S2E, S2G, S4B, S6F, S7C) were obtained by first segmenting the cell using ‘Threshold’ option of Fiji/ImageJ to obtain a binary image which properly covered all pixels of the cell. Subsequently, using the “Analyze Particle” option, cell masks were generated. For optimizing these masks, we used ‘Fill holes’, ‘Erode’ and ‘Dilate’ functions. Finally, the ‘Outline’ command was operated on binarized cells, and ‘Temporal-Color Code’ was used^4,35^.

For membrane kymographs (Figures 3I-J, 3M-N, 5F, 6F, 7D, 7M, S4C, S6G, S7D), cells were segmented against the background following standard image-processing steps with custom code written in MATLAB^4,35,88^. Next, kymographs were created from the segmented cells by aligning consecutive lines over time by minimizing the sum of the Euclidean distances between the coordinates in two consecutive frames using a custom-written MATLAB function^65^. A linear colourmap was used for the normalized intensities in the kymographs; the lowest intensity is indicated by blue and the highest with yellow.

For linescan analysis (Figure 3H, L), a straight-line segment (width of 5 pixels) was drawn, using the ‘Straight’ tool in Fiji/ImageJ, across the cellular protrusion region to obtain an average intensity value. The intensity values along that particular line were obtained using the ‘Plot Profile’ option. These values were plotted along the distance of the line in Microsoft Excel^35,88^.

Local protrusion (Figure 2D, F, L) and cell migration (Figures 5G-K, 6G-K, 7E-I, 7N-R, S6H-L, S7E-I) analyses were performed as described previously^4,35,88^. For cell speed, area and aspect ratio quantifications (in Figures 3D-F, 4C-E, 4G-H), cells were segmented against the background following standard image-processing steps with custom code written in Python based on package scikit-image, Trackpy and PyImageJ. Human supervision was involved during the process with an integration to Fiji, and we obtained the X-position, Y-position, cell area, and aspect ratio for each frame of each cell. Distance between adjacent frames was calculated from X-position and Y-position, and was divided by time interval to obtain an instantaneous velocity. The average speed of cells was calculated by taking an average of all instantaneous velocities. For Figure S1B, macropinocytosis uptake was analyzed by outlining each cell and quantifying the total TRITC fluorescence signal within each outline divided by the cell area^89^. For Figure S4B, numbers of blebs for each cell, before or after recruitment, were quantified by summing them over a min^96^.

### Simulations

The simulations are based on a model, previously described, in which three interacting species, RasGTP, PIP2, and PKB, form an excitable network^37^. To simplify the notation, we describe the concentration of these three species using the symbols: *F*, *B* and *R*, respectively, which represent the fact that RasGTP and PIP2 are front- and back-associated molecules, respectively, and that PKB acts to provide the refractory element of the excitable network behavior.

The concentrations of each of these molecules is described by stochastic, reaction-diffusion partial differential equations:

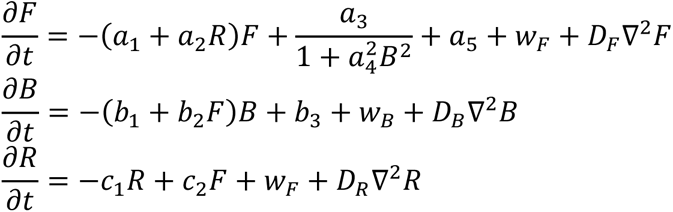

In each of these equations, the final term represents the diffusion of the species, where *D*_∗_ is the respective diffusion coefficient and ∇^2^ is the spatial Laplacian (in one or two dimensions). The second-to-last terms represent the molecular noise. Our model assumes a Langevin approximation in which the size of the noise is based on the reaction terms^97^. For example, in the case of PKB (*R*), the noise is given by

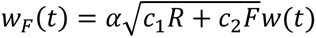

where *w*(*t*) is a zero mean, unit variance Gaussian Brown noise process. In the simulations, the size of this noise was adjusted with the empirical parameter α.

In addition to the EN dynamics described above, we incorporated two other terms related to cell polarization^37,62^. These feedback loops come from *B* and *R*:

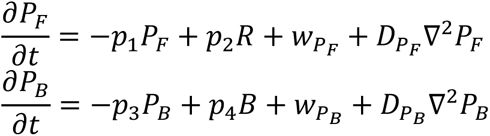

They modify the equation for Ras:

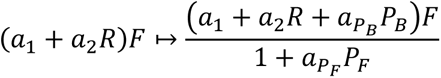

having the effect of increasing front and back contributions. Lastly, we note that changes in RasGAP contributions are modeled as changes in the parameters *a*_1_ and *a*_2_.

Simulations were run on MATLAB 2023a (Mathworks, Natick, MA) on custom-code based on the Ito solution in the Stochastic Differential Equation toolbox (http://sdetoolbox.sourceforge.net). Two-dimensional simulations were used to recreate the observed wave patterns of larger electrofused cells, and so assume a grid 40 µm×40 µm with a spacing of 0.4 µm × 0.4 µm µm per grid point (i.e., 100× 100 points) and zero flux boundary conditions. The one-dimensional simulations aim to recreate the membrane fluorescence observed in single-cell confocal images. The dimension is therefore smaller, assuming a cell radius of 5 µm and a spacing of 0.25 µm, resulting in 2π × 5/0.25 ≈ 126 points along the perimeter, and periodic boundary conditions.

Parameter values

| Parameter | Value | Units | Parameter | Value | Units |
| --- | --- | --- | --- | --- | --- |
| $a_1$ | 1 | $\text{s}^{-1}$ | $b_1$ | 0.06 | $\text{s}^{-1}$ |
| $a_2$ | 37 | $\mu\text{M}^{-1} \text{s}^{-1}$ | $b_2$ | 8880 | $\mu\text{M}^{-1} \text{s}^{-1}$ |
| $a_3$ | 12 | $\text{s}^{-1}$ | $b_3$ | 8.20 | $\mu\text{M} \text{s}^{-1}$ |
| $a_4$ | 444 | $\mu\text{M}^{-1}$ | $c_1$ | 0.15 | $\mu\text{M}^{-1} \text{s}^{-1}$ |
| $a_5$ | $5.4 \times 10^{-3}$ | $\text{s}^{-1}$ | $c_2 (1D)$ | 1.50 | $\mu\text{M}^{-1} \text{s}^{-1}$ |
| $D_F$ | 0.15 | $\mu\text{m}^2/\text{s}$ | $c_2 (2D)$ | 0.75 | $\mu\text{M}^{-1} \text{s}^{-1}$ |
| $D_B$ | 0.15 | $\mu\text{m}^2/\text{s}$ | $p_1$ | 0.05 | $\mu\text{M}^{-1} \text{s}^{-1}$ |
| $p_2$ | 0.05 | $\mu\text{M}^{-1} \text{s}^{-1}$ | $p_3$ | 0.05 | $\mu\text{M}^{-1} \text{s}^{-1}$ |
| $p_4$ | 0.05 | $\mu\text{M}^{-1} \text{s}^{-1}$ | $D_R (1D)$ | 1.50 | $\mu\text{m}^2/\text{s}$ |
| $D_R (2D)$ | 0.25 | $\mu\text{m}^2/\text{s}$ | $D_{P_F}$ | 0.0125 | $\mu\text{m}^2/\text{s}$ |
| $D_{P_B}$ | 0.0125 | $\mu\text{m}^2/\text{s}$ | $a_{P_F}$ | 2 | $\mu\text{M}^{-1}$ |
| $a_{P_B}$ | 1 | $\mu\text{M}^{-1} \text{s}^{-1}$ | $\alpha$ (1D) | 0.0833 | |
| $\alpha$ (2D) | 0.0417 | | | | |

### Statistical analysis

Statistical analyses were executed using unpaired or paired 2-tailed non-parametric tests on GraphPad Prism 8. Results are expressed as mean ± SD from at least 3 independent experiments. ns denotes P>0.05, * denotes P ≤ 0.05, ** denotes P ≤ 0.01, *** denotes P ≤ 0.001, **** denotes P ≤ 0.0001.

## Data availability

All data are provided in the main or supplementary text. Requests for additional information on this work are to be made to the corresponding authors.

## Supplementary Information

### Supplementary Figures

**Figure S1.**
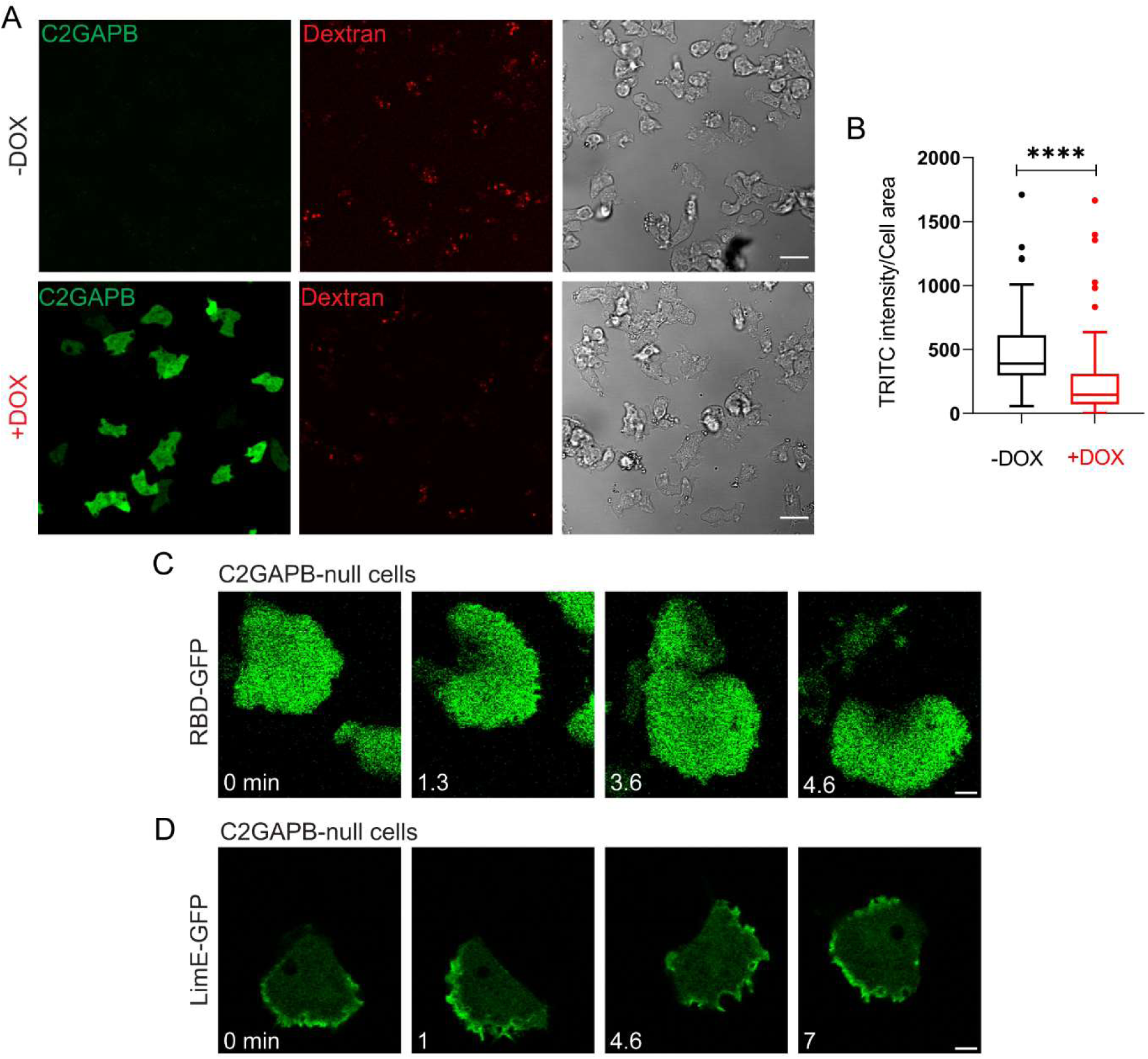
C2GAPB regulates Ras and protrusive activities. **(A)** Representative confocal images of vegetative *Dictyostelium* cells before (top panel, ‘-DOX’) and after (bottom panel, ‘+DOX’) doxycycline-induced GFP-C2GAPB (green) expression. Cells (DIC) were treated with TRITC-dextran (red) before imaging. Scale bars represent 10 µm. **(B)** Quantification of macropinocytosis uptake, before (black) and after (red) C2GAPB expression. n_c_=103 from atleast 3 independent experiments; asterisks indicate significant difference, ****P ≤ 0.0001 (Wilcoxon-Mann-Whitney rank sum test). **(C, D)** Time-lapse confocal images of vegetative C2GAPB-null *Dictyostelium* cells expressing **(C)** RBD-GFP or **(D)** LimE-GFP. Time in ‘min’ format. Scale bars represent 5 µm.

**Figure S2.**
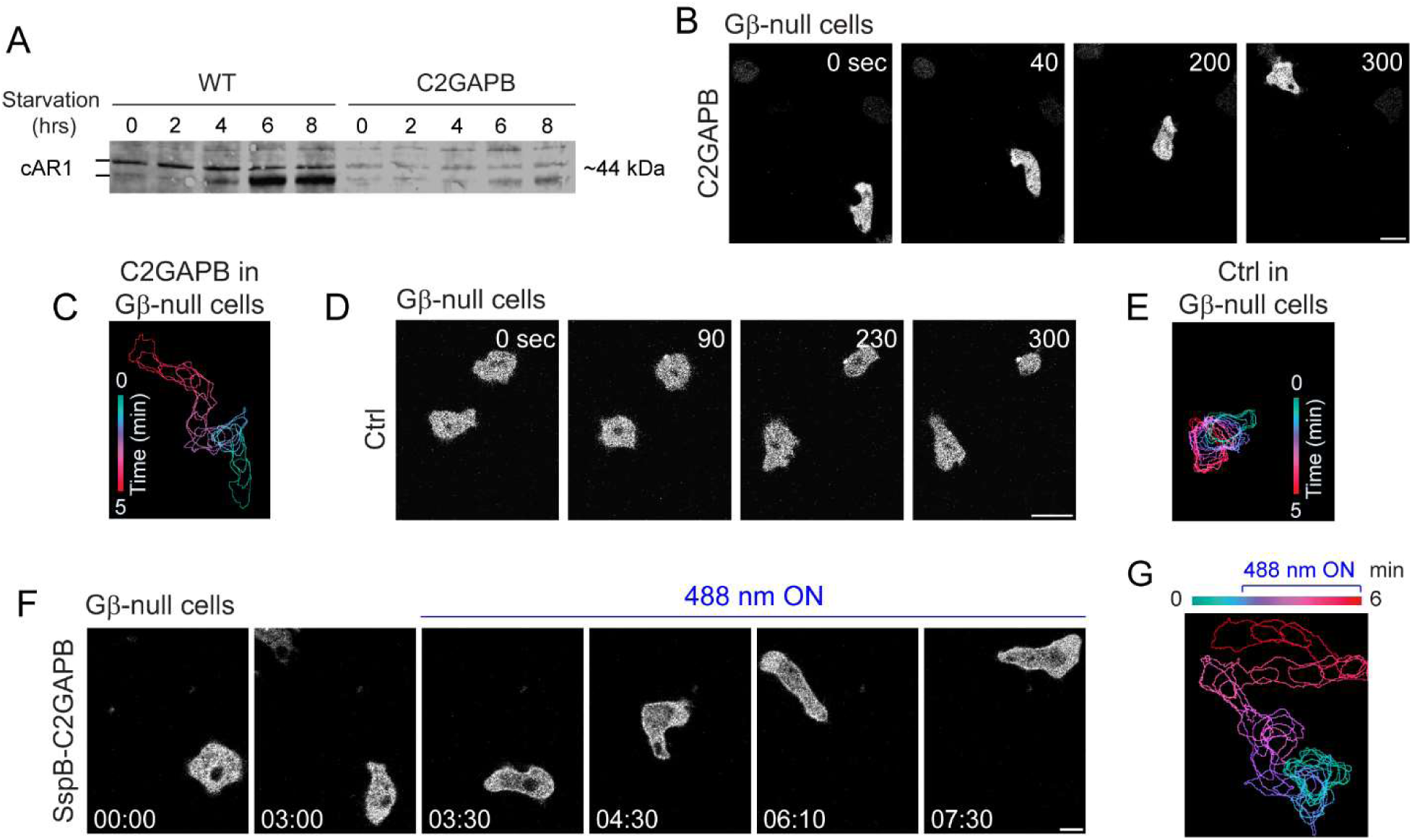
C2GAPB-induced polarization does not require development or GPCR signaling. **(A)** Representative western blot showing expression of cAR1 (∼44 kDa) in developing wildtype (WT) or C2GAPB-expressing *Dictyostelium* cells during starvation (0-8 hrs). cAR1 appears as a doublet denoting its two forms, unmodified (lower band) and phosphorylated (upper band). Time-lapse confocal images of vegetative Gβ-null *Dictyostelium* cells expressing **(B)** mRFPmars-C2GAPB or **(D)** tgRFPt-Ctrl (control without C2GAPB) after overnight doxycycline treatment. Time in ‘sec’ format. Scale bars represent 5 µm. **(C, E)** Color-coded (at 1-min interval) outlines of the C2GAPB- or Ctrl-expressing cell shown in (B) or (D), respectively. **(F)** Time-lapse confocal images of Gβ-null *Dictyostelium* cell expressing mRFPmars-SspB R73Q-C2GAPB, before or after 488 nm laser was switched on globally. Time in min:sec format. Scale bars represent 5 µm. **(G)** Color-coded (at 1-min interval) outlines of the mRFPmars-SspB R73Q-C2GAPB-expressing cell shown in (F).

**Figure S3.**
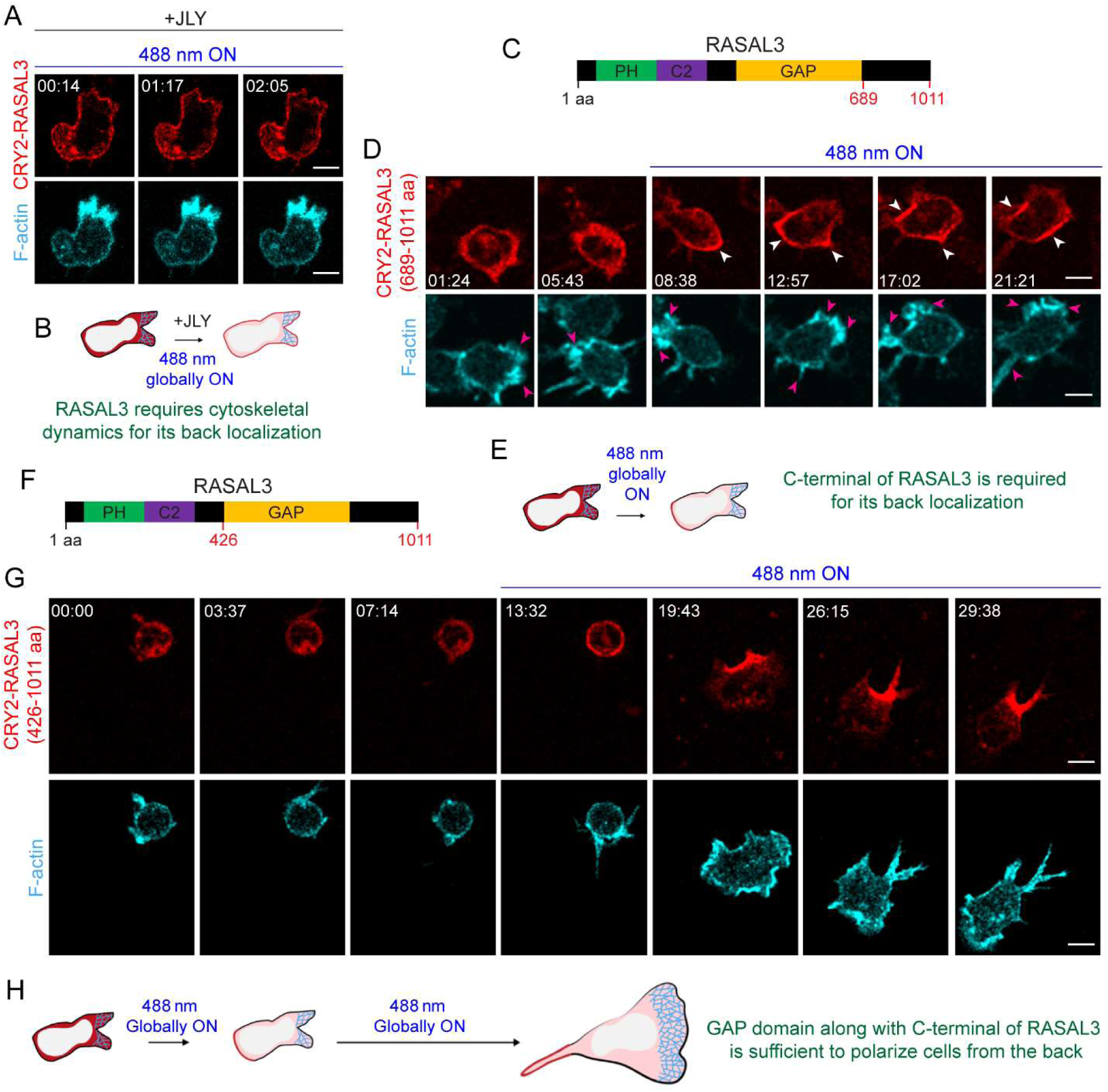
RASAL3 back localization is dependent on cytoskeletal dynamics and its C-terminal tail whereas polarizing function of RASAL3 is due to its GAP domain along with C-terminal tail. **(A)** Time-lapse confocal images of JLY cocktail-treated HL-60 neutrophil expressing CRY2PHR-mCherry-RASAL3 (red; upper panel) and LifeAct-miRFP703 (cyan; lower panel), after 488 nm laser was turned on globally. Time in min:sec format. Scale bars represent 5 µm. **(B)** Cartoon shows that JLY treatment caused RASAL3 to recruit uniformly instead of localizing to the back. **(C, F)** Schematics showing RASAL3 protein sequence (1-1011 amino acid long). Two truncation mutants were generated in this study: **(C)** RASAL3_689-1011_ which consists of only the C-terminal tail (689-1011 amino acids) and **(F)** RASAL3_426-1011_ consisting of the GAP domain along with the C-terminal tail (426-1011 amino acids). **(D, G)** Time-lapse confocal images of differentiated HL-60 neutrophil expressing **(D)** CRY2PHR-mCherry-RASAL3 (689-1011 aa) or **(G)** CRY2PHR-mCherry-RASAL3 (426-1011 aa) (red; upper panel) and LifeAct-miRFP703 (cyan; lower panel), before or after 488 nm laser was turned on globally. Time in min:sec format. Scale bars represent 5 µm. **(E, H)** Cartoons demonstrate phenomenon observed with recruiting CRY2PHR-mCherry-RASAL3 (689-1011 aa) or CRY2PHR-mCherry-RASAL3 (426-1011 aa) in differentiated neutrophils.

**Figure S4.**
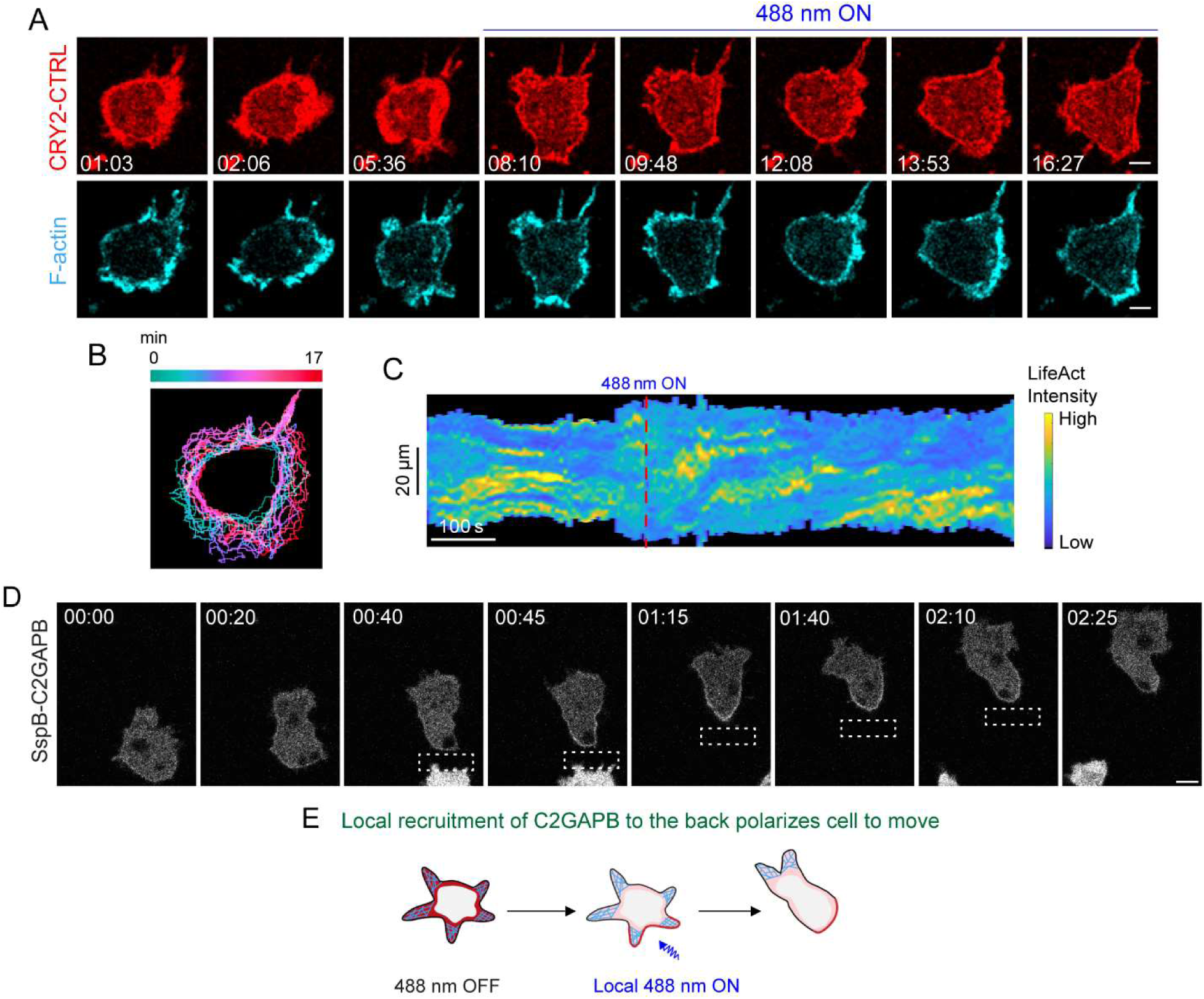
Recruitment of RasGAP, and not CTRL, moves cell rapidly. **(A)** Time-lapse confocal images of differentiated HL-60 neutrophil expressing CRY2PHR-mCherry-CTRL (control without RASAL3, red; upper panel) and LifeAct-miRFP703 (cyan; lower panel), before or after 488 nm laser was globally applied. Time in min:sec format. Scale bars represent 5 µm. **(B)** Color-coded (at 1 min intervals) outlines of the cell shown in (A). **(C)** Representative kymograph of cortical LifeAct intensity in CTRL-expressing neutrophil before or after 488 nm laser was switched on. A linear color map shows that blue is the lowest LifeAct intensity whereas yellow is the highest. Duration of the kymograph is 17 mins. The cartoon summarizes membrane recruitment, F-actin polymerization or cell shape status corresponding to the kymograph. **(D)** Time-lapse confocal images of vegetative *Dictyostelium* cells expressing mRFPmars-SspB R73Q-C2GAPB which is recruited to the back of the migrating cell by applying 488 nm laser near it, as shown by the dashed white box. Time in min:sec format. Scale bars represent 5 µm. **(E)** Cartoon demonstrates C2GAPB-mediated phenomenon shown in (D).

**Figure S5.**
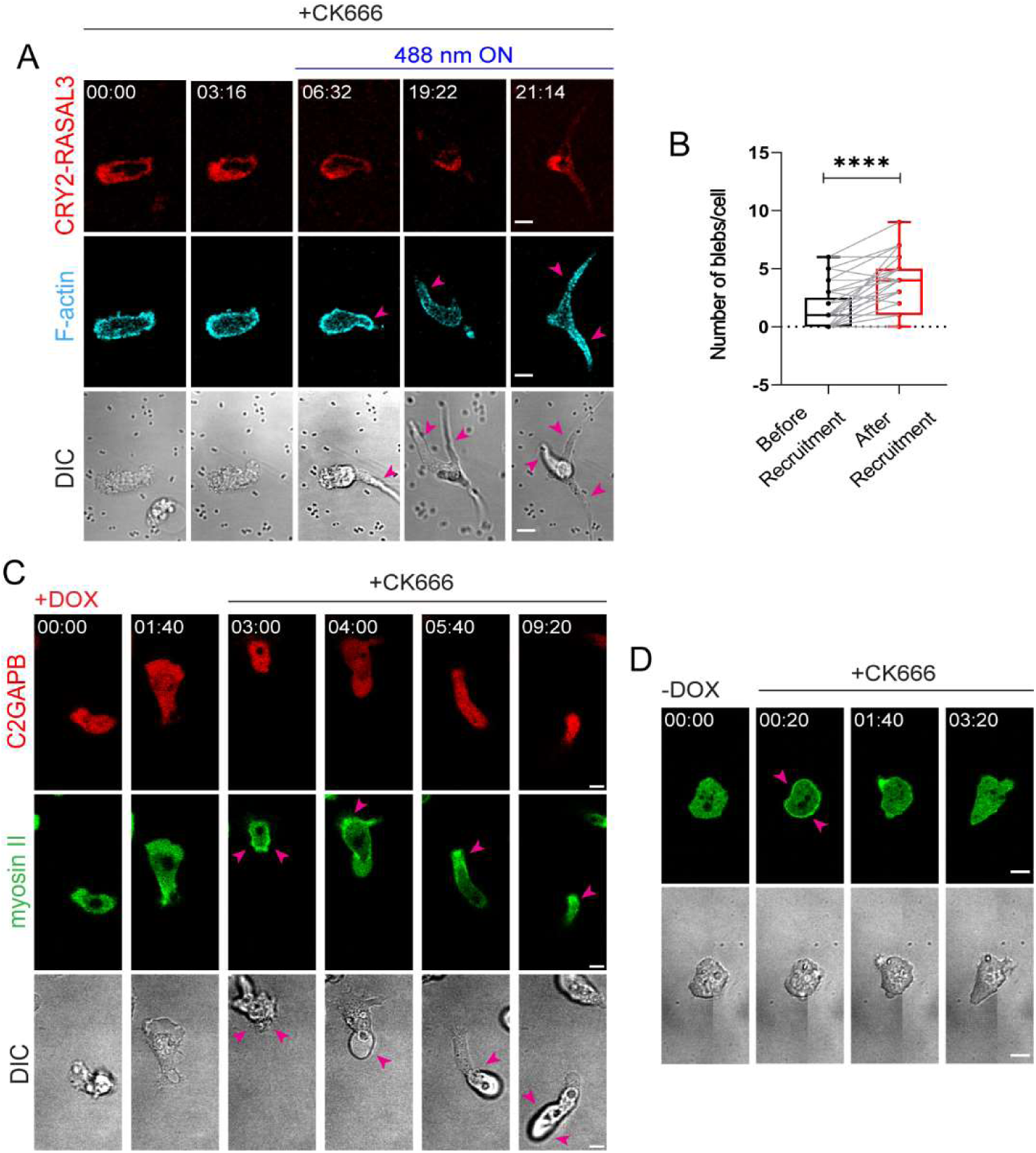
RasGAPs increase cellular contraction. **(A)** Time-lapse confocal images of CK666-treated differentiated HL-60 neutrophil expressing CRY2PHR-mCherry-RASAL3 (red; upper panel) and LifeAct-miRFP703 (cyan; lower panel), before or after 488 nm laser was globally applied. Pink arrows denote long blebs which appear after recruited RASAL3 localized to the back of CK666-treated cell. Time in min:sec format. Scale bars represent 5 µm. **(B)** Box-and-whisker plot of number of blebs per cell within a minute, before (black) or after (red) RASAL3 recruitment. n_c_=37 from atleast 3 independent experiments; asterisks indicate significant difference, ****P ≤ 0.0001 (Wilcoxon-Mann-Whitney rank sum test). Time-lapse confocal images of vegetative *Dictyostelium* cells expressing mRFPmars-C2GAPB (red; top panel) and myosin II-GFP (green; middle panel) before **(C)** or after **(D)** overnight doxycycline treatment. CK666 was added during imaging. Since there is no C2GAPB expression without doxycycline treatment, C2GAPB (red panel) is not shown in (D). Pink arrows denote appearance of long blebs (DIC channel) in C2GAPB-expressing cells after CK666 was added. Time in min:sec format. Scale bars represent 5 µm.

**Figure S6.**
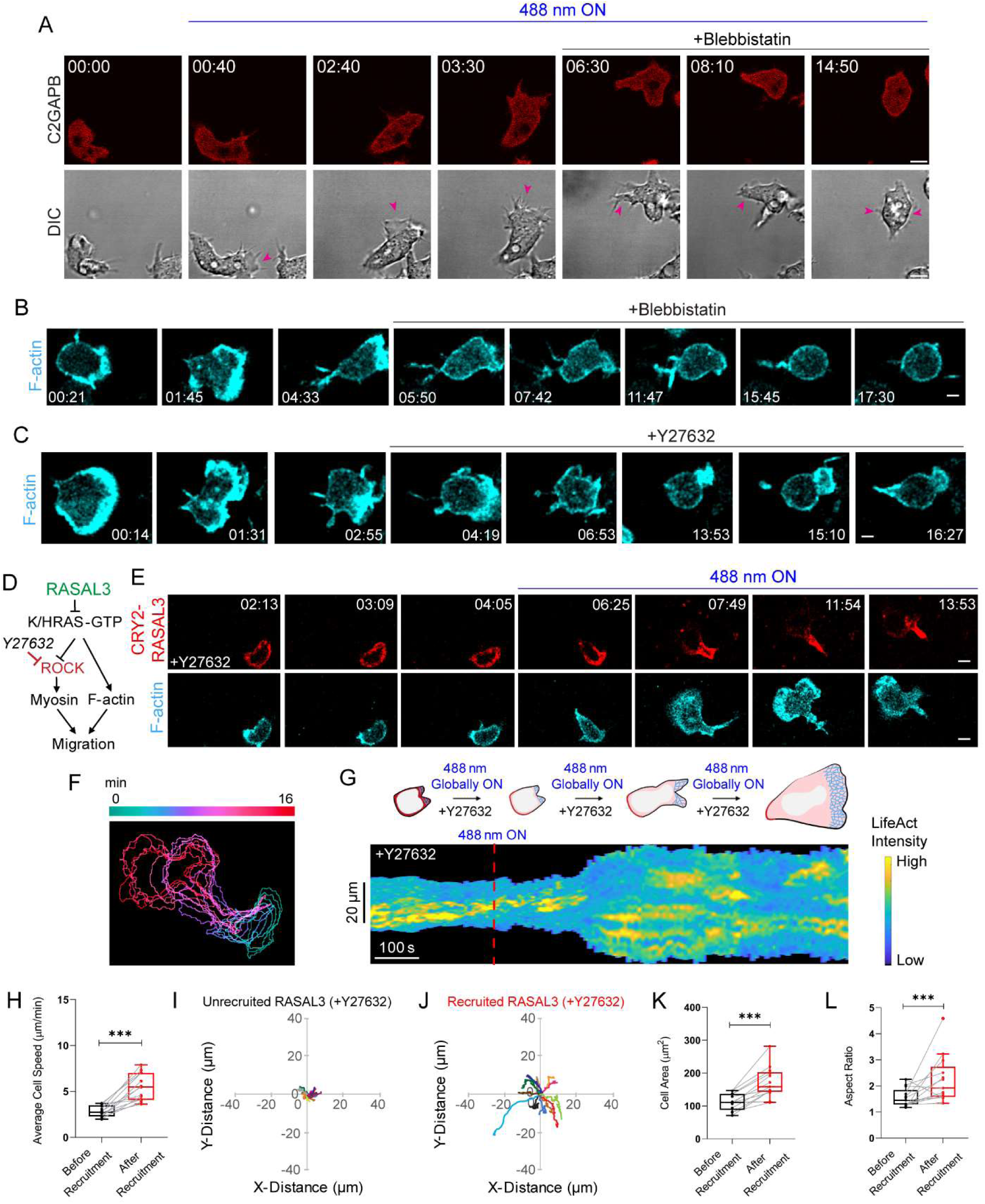
Global RASAL3 recruitment overcomes ROCK inhibition whereas global C2GAPB recruitment does not overcome myosin inhibition. **(A)** Time-lapse confocal images of vegetative *Dictyostelium* cells expressing mRFPmars-SspB R73Q-C2GAPB (red; top panel), before or after 488 nm laser was switched on globally. Blebbistatin was added during imaging. Pink arrows denote cell of interest (DIC channel). Time in min:sec format. Scale bars represent 5 µm. **(B, C)** Time-lapse confocal images of HL-60 neutrophil expressing LifeAct-miRFP703 (cyan; lower panel), before or after **(B)** blebbistatin and **(C)** Y27632 treatment. Time in min:sec format. Scale bars represent 5 µm. **(D)** Strategy for testing effect of ROCK inhibitor, Y27632, on RASAL3-directed actin polymerization and migration. **(E)** Time-lapse confocal images of Y27632-treated HL-60 neutrophil expressing CRY2PHR-mCherry-RASAL3 (red; upper panel) and LifeAct-miRFP703 (cyan; lower panel), before or after 488 nm laser was turned on globally. Time in min:sec format. Scale bars represent 5 µm. **(F)** Color-coded (at 1 min intervals) outlines of the cell shown in (E). **(G)** Representative kymograph of cortical LifeAct intensity in Y27632-treated RASAL3-expressing neutrophil before or after 488 nm laser was turned on. A linear color map shows that blue is the lowest LifeAct intensity whereas yellow is the highest. Duration of the kymograph is 16 mins. Cartoon summarizes membrane recruitment, actin polymerization or cell shape status corresponding to the kymograph. Box-and-whisker plots of **(H)** cell speed, **(K)** cell area, and **(L)** aspect ratio, before (black) and after (red) RASAL3 recruitment in Y27632-treated cells. n_c_=13 from atleast 3 independent experiments; asterisks indicate significant difference, ***P ≤ 0.001 (Wilcoxon-Mann-Whitney rank sum test). Centroid tracks of Y27632-treated neutrophils (n_c_=13) showing random motility before **(I)** or after **(J)** RASAL3 recruitment. Each track lasts atleast 5 mins and was reset to same origin.

**Figure S7.**
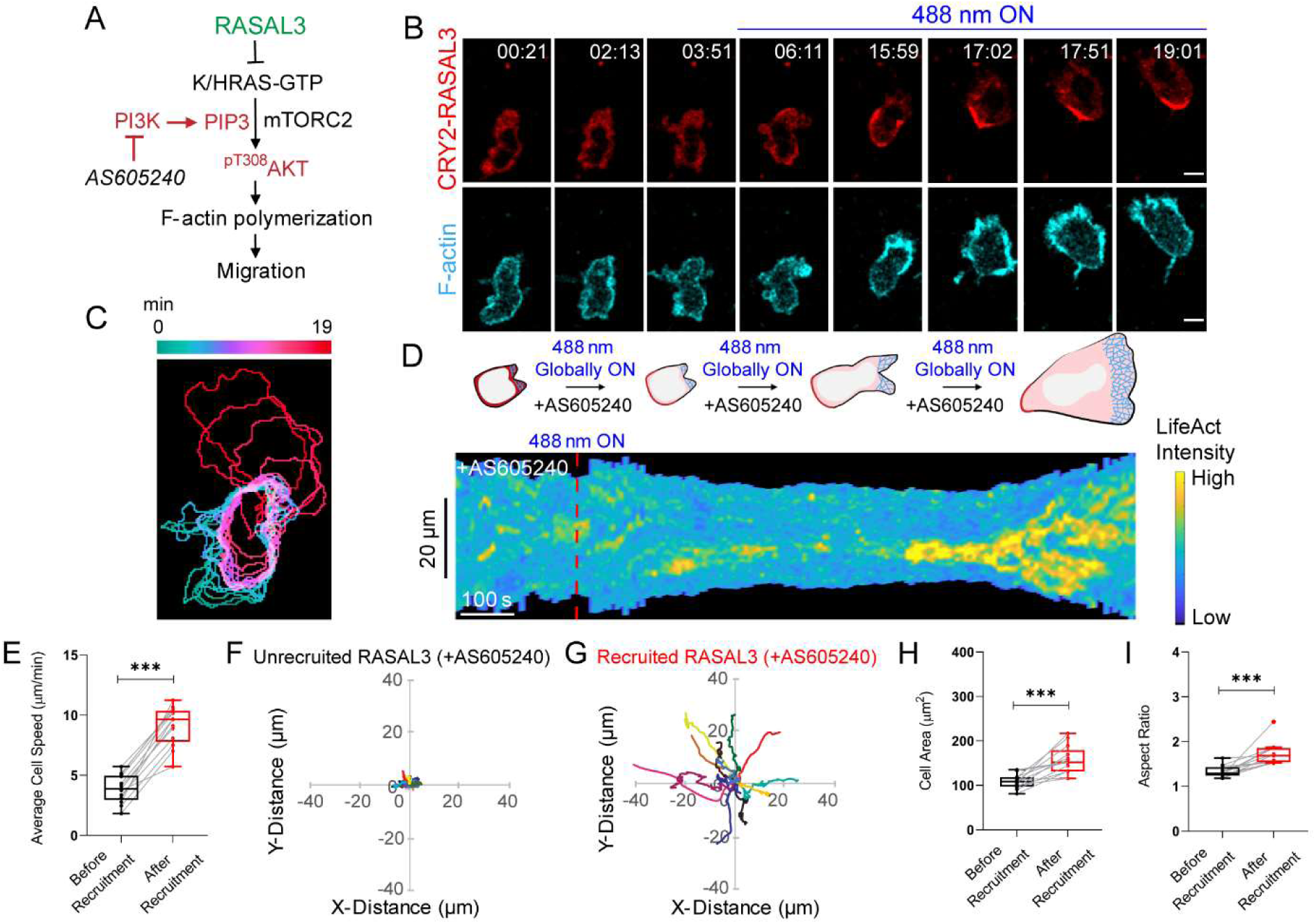
Global RASAL3 recruitment overcomes PI3Kγ inhibition. **(A)** Strategy for testing effect of PI3Kγ inhibitor, AS605240, on RASAL3-directed actin polymerization and motility. **(B)** Time-lapse confocal images of AS605240-treated HL-60 neutrophil expressing CRY2PHR-mCherry-RASAL3 (red; upper panel) and LifeAct-miRFP703 (cyan; lower panel), before or after 488 nm laser was turned on globally. Time in min:sec format. Scale bars represent 5 µm. **(C)** Color-coded (at 1 min intervals) outlines of the cells shown in (B). **(D)** Representative kymograph of cortical LifeAct intensity in AS605240-treated RASAL3-expressing neutrophil before or after 488 nm laser was turned on. A linear color map shows that blue is the lowest LifeAct intensity whereas yellow is the highest. Duration of the kymograph is 19 mins. Cartoon summarizes membrane recruitment, actin polymerization or cell shape status corresponding to the kymograph. Box-and-whisker plots of **(E)** cell speed, **(H)** cell area, and **(I)** aspect ratio, before (black) and after (red) RASAL3 recruitment in AS605240-treated cells. n_c_=13 from atleast 3 independent experiments; asterisks indicate significant difference, ***P ≤ 0.001 (Wilcoxon-Mann-Whitney rank sum test). Centroid tracks of AS605240-treated neutrophils (n_c_=13) showing random motility before **(F)** or after **(G)** RASAL3 recruitment. Each track lasts atleast 5 mins and was reset to same origin.

## Supplementary Video Legends

**Video S1**

Time-lapse confocal microscopy of GFP-RBD (green; left panel) cortical waves at the substrate-attached surface in electrofused, giant *Dictyostelium* cell, before overnight doxycycline treatment (‘-DOX’ at top of the video). Without doxycycline, mRFPmars-C2GAPB (red; right panel) was not expressed in the cell. Top left corner shows time in min:sec format. Scale bar represents 5 µm.

**Video S2**

Time-lapse confocal microscopy of GFP-RBD (green; left panel) cortical waves at the substrate-attached surface in electrofused, giant *Dictyostelium* cells, after overnight doxycycline treatment (‘+DOX’ at top of the video). Doxycycline induced mRFPmars-C2GAPB (red; right panel) expression in the cell. Top left corner shows time in min:sec format. Scale bar represents 5 µm.

**Video S3**

Time-lapse confocal microscopy of PHcrac-YFP (green; left panel) cortical waves at the substrate-attached surface in electrofused, giant *Dictyostelium* cell, before overnight doxycycline treatment (‘-DOX’ at top of the video). Without doxycycline, mRFPmars-C2GAPB (red; right panel) was not expressed in the cell. Top left corner shows time in min:sec format. Scale bar represents 5 µm.

**Video S4**

Time-lapse confocal microscopy of PHcrac-YFP (green; left panel) cortical waves at the substrate-attached surface in electrofused, giant *Dictyostelium* cells, after overnight doxycycline treatment (‘+DOX’ at top of the video). Doxycycline induced mRFPmars-C2GAPB (red; right panel) expression in the cell. Top left corner shows the time in min:sec format. Scale bar represents 5 µm.

**Video S5**

Time-lapse confocal microscopy of RBD-YFP (yellow; left panel) cortical waves at the substrate-attached surface of electrofused, giant *Dictyostelium* cells, before or after global membrane recruitment of mRFPmars-SspB R73Q-C2GAPB (‘Opto-C2GAPB’; red, right panel). Global recruitment was initiated when blue laser was applied globally (“488nm on” at top of the video) at ‘05:50’ and ‘30:00’. Propagating RBD waves extinguished with C2GAPB recruitment, and recovered when 488nm laser was switched off again. Top left corner shows the time in min:sec format. Scale bars represent 10 µm.

**Video S6**

Time-lapse confocal microscopy of PHcrac-YFP (yellow; left panel) cortical waves at the substrate-attached surface of electrofused, giant *Dictyostelium* cells, before or after global membrane recruitment of mRFPmars-SspB R73Q-C2GAPB (‘Opto-C2GAPB’; red, right panel). Global recruitment was initiated when blue laser was applied globally (“488nm on” at top of the video) at ‘16:20’ or ‘02:00’ in movie 1 or 2, respectively. Propagating PHcrac waves extinguished with C2GAPB recruitment. Top left corner shows the time in min:sec format. Scale bars represent 10 µm.

**Video S7**

Time-lapse confocal microscopy of differentiated HL-60 neutrophil expressing CRY2PHR-mCherry-RASAL3 (red; left panel) and LifeAct-miRFP703 (cyan; right panel). RASAL3 was recruited to the membrane anchor, CIBN-CAAX (untagged), at the cell front by intermittently applying blue (488 nm) laser near it, as shown with the white box in the red channel. The disappearance of cellular protrusions at the RASAL3 recruitment site is denoted with solid white box in the blue channel. The top right corner shows time in min:sec format. Neutrophils were not exposed to chemoattractants during this experiment. Scale bar: 5 µm.

**Video S8**

Time-lapse confocal microscopy of vegetative *Dictyostelium* AX2 cells before (-Dox; left panel) and after overnight doxycycline treatment (+Dox; right panel). Doxycycline induced mRFPmars-C2GAPB (red; right panel) expression in cells. The top right corner of each panel shows time in min:sec format. Cells were not exposed to chemoattractants during this experiment. Scale bar represents 20 µm.

**Video S9**

Time-lapse confocal microscopy of vegetative *Dictyostelium* AX2 cell before or after global membrane recruitment of tgRFPt-SspB R73Q-Ctrl (Opto-Ctrl). Global recruitment was initiated when blue laser was applied globally (“488nm on” at top of the video) at ‘04:00’. Top left corner shows time in min:sec format. Cell was not exposed to any chemoattractant during this experiment. Scale bar represents 5 µm.

**Video S10**

Time-lapse confocal microscopy of vegetative *Dictyostelium* AX2 cell before or after global membrane recruitment of mRFPmars-SspB R73Q-C2GAPB (Opto-C2GAPB). In movie 1, global recruitment was initiated when blue laser was applied globally (“488nm on” at top of the video) at ‘02:00’. In movie 2, blue laser was switched on or off multiple times during the course of the experiment, as denoted with “488nm on” or “488nm off” at the top of the movie. Top left corner shows time in min:sec format. Cells were not exposed to any chemoattractant during this experiment. Scale bar represents 5 µm.

**Video S11**

Time-lapse confocal microscopy of vegetative *Dictyostelium* Gβ null (Gβ^-^) cell before or after global membrane recruitment of mRFPmars-SspB R73Q-C2GAPB (Opto-C2GAPB). Global recruitment was initiated when blue laser was applied globally (“488nm on” at top of the video) at ‘08:20’. Top left corner shows time in min:sec format. Cell was not exposed to any chemoattractant during this experiment. Scale bar represents 5 µm.

**Video S12**

Time-lapse confocal microscopy of LimE_Δcoil_-YFP (yellow; left panel) cortical waves at the substrate-attached surface of electrofused, giant *Dictyostelium* cells, before or after global membrane recruitment of mRFPmars-SspB R73Q-C2GAPB (‘Opto-C2GAPB’; red, right panel). Global recruitment was initiated when blue laser was applied globally (“488nm on” at top of the video) at ‘15:00’. Propagating LimE waves extinguished, with C2GAPB recruitment, except for one standing wave in the middle. Top left corner shows the time in min:sec format. Scale bar represents 10 µm.

**Video S13**

Time-lapse confocal microscopy of differentiated, unpolarized HL-60 neutrophil expressing CRY2PHR-mCherry-RASAL3 (red; left panel) and LifeAct-miRFP703 (cyan; right panel). RASAL3 was recruited to the membrane anchor, CIBN-CAAX (untagged), at the transient protrusions by intermittently applying blue (488 nm) laser near it, as shown with the white box in the red channel. The disappearance of cellular protrusions at the RASAL3 recruitment site is denoted with solid white box in the blue channel. 488 nm laser was switched off ‘03:09’ onwards to visualize the effects of RASAL3 self-rearrangement on the membrane. The top right corner shows time in min:sec format. Neutrophils were not exposed to chemoattractants during this experiment. Scale bar: 5 µm.

**Video S14**

Time-lapse confocal microscopy of differentiated HL-60 neutrophil expressing CRY2PHR-mCherry-RASAL3 (red; left panel) and LifeAct-miRFP703 (cyan; right panel), before or after 488 nm laser was applied globally. The membrane anchor, untagged CIBN-CAAX, was expressed. Top left corner shows time in min:sec format. To start recruitment (red; left panel), the laser was switched on at ‘07:07’ once ‘488 nm ON’ appears at the top of the video. Cell was not exposed to chemoattractant during the experiment. Scale bar represents 5 µm.

**Video S15**

Time-lapse confocal microscopy of differentiated HL-60 neutrophil expressing CRY2PHR-mCherry-CTRL (red; left panel) and LifeAct-miRFP703 (cyan; right panel), before or after 488 nm laser was turned on globally. Membrane anchor, untagged CIBN-CAAX, was also expressed. Top left corner shows time in min:sec format. To start recruitment (red; left panel), the laser was switched on at ‘05:43’ once ‘488 nm ON’ appears at the top of the video. Cell was not exposed to chemoattractant during the experiment. Scale bar represents 5 µm.

**Video S16**

Time-lapse confocal microscopy of differentiated, blebbistatin-treated HL-60 neutrophil expressing CRY2PHR-mCherry-RASAL3 (red; left panel) and LifeAct-miRFP703 (cyan; right panel), before or after 488 nm laser was turned on globally. Membrane anchor, untagged CIBN-CAAX, was also expressed. Cell was treated with 20 µM blebbistatin atleast 10 mins before imaging. Pink arrow denotes the cell of interest. To start recruitment (red; left panel), the laser was switched on at ‘03:23’ once ‘488 nm ON’ appears at the top of the video. Cell was not exposed to chemoattractant during the experiment. Top left corner shows time in min:sec format. Scale bar represents 5 µm.

**Video S17**

Time-lapse confocal microscopy of differentiated, Y27632-treated HL-60 neutrophil expressing CRY2PHR-mCherry-RASAL3 (red; left panel) and LifeAct-miRFP703 (cyan; right panel), before or after 488 nm laser was turned on globally. Membrane anchor, untagged CIBN-CAAX, was also expressed. Cell was treated with 10 µM Y27632 atleast 10 mins before imaging. To start recruitment (red; left panel), the laser was switched on at ‘04:12’ once ‘488 nm ON’ appears at the top of the video. Neutrophil was not exposed to any chemoattractant during imaging. Top left corner shows time in min:sec format. Scale bar represents 5 µm.

**Video S18**

Time-lapse confocal microscopy of differentiated, LY294002-treated HL-60 neutrophil expressing CRY2PHR-mCherry-RASAL3 (red; left panel) and LifeAct-miRFP703 (cyan; right panel), before or after 488 nm laser was applied globally. The membrane anchor, untagged CIBN-CAAX, was expressed as well. Cell was treated with 50 µM LY294002 atleast 10 mins before imaging. To start recruitment (red; left panel), the laser was switched on at ‘04:12’ once ‘488 nm ON’ appears at the top of the video. Neutrophil was not exposed to any chemoattractant during imaging. Top left corner shows time in min:sec format. Scale bar represents 5 µm.

**Video S19**

Time-lapse confocal microscopy of differentiated, AS605240-treated HL-60 neutrophil expressing CRY2PHR-mCherry-RASAL3 (red; left panel) and LifeAct-miRFP703 (cyan; right panel), before or after 488 nm laser was turned on globally. Membrane anchor, untagged CIBN-CAAX, was expressed here. Cell was treated with 20 µM AS605240 atleast 10 mins before imaging. To start recruitment (red; left panel), the laser was turned on at ‘03:58’ once ‘488 nm ON’ appears at the top of the video. Neutrophil was not exposed to any chemoattractant during imaging. Top left corner shows time in min:sec format. Scale bar represents 5 µm.

**Video S20**

Time-lapse confocal microscopy of differentiated, PP242-treated HL-60 neutrophil expressing CRY2PHR-mCherry-RASAL3 (red; left panel) and LifeAct-miRFP703 (cyan; right panel), before or after 488 nm laser was turned on globally. Membrane anchor, untagged CIBN-CAAX, was expressed. Cell was treated with 20 µM PP242, 10 mins before imaging. To start recruitment (red; left panel), the laser was turned on at ‘06:04’ once ‘488 nm ON’ appears at the top of the video. Cell was not exposed to any chemoattractant during imaging. Top left corner shows time in min:sec format. Scale bar represents 5 µm.

**Video S21**

Two-dimensional simulation results of the excitable network for varying RasGAP levels. The video shows three different simulations, with 80% WT, WT and 110%-RasGAP levels, sequentially. The area is a square with sides 40 μm long and the time-stamp denotes seconds. The three colors correspond to Ras (red), PIP2 (green) and PKB (blue).

## Notes

### Competing Interest Statement

The authors have declared no competing interest.

